# SHAPE-based chemical probes for studying preQ_1_-RNA interactions in living bacteria

**DOI:** 10.1101/2025.07.21.665968

**Authors:** José A. Reyes Franceschi, Emilio L. Cárdenas, Brandon J. C. Klein, Chase A. Weidmann, Amanda L. Garner

**Author notes:** These authors contributed equally to this work (José A. Reyes Franceschi, Emilio L. Cárdenas).

## Abstract

Interrogating RNA-small molecule interactions inside cells is critical for advancing RNA-targeted drug discovery. In particular, chemical probing technologies that both identify small molecule-bound RNAs and define their binding sites in the complex cellular environment will be key for establishing the on-target activity necessary for successful hit-to-lead campaigns. Using the small molecule metabolite preQ_1_ and its cognate riboswitch RNA as a model, herein we describe a chemical probing strategy for filling this technological gap. Building on well-established RNA acylation chemistry employed by *in vivo* click selective 2′-hydroxyl acylation analyzed by primer extension (icSHAPE) probes, we developed an icSHAPE-based preQ_1_ probe that retains biological activity in a preQ_1_ riboswitch reporter assay and successfully enriches the preQ_1_ riboswitch from living bacterial cells. Further, we map the preQ_1_ binding site on probe-modified riboswitch RNA by mutational profiling (MaP). As the need for rapid profiling of on- and off-target small molecule interactions continues to grow, this chemical probing strategy offers a method to interrogate cellular RNA-small molecule interactions and support the future development of RNA-targeted therapeutics.

## INTRODUCTION

Cellular RNAs fold into a complex array of secondary and tertiary structures to carry out their functions.^1,2^ Relevant to chemical biology and drug discovery, many of these structures are predicted to bind small molecules, and many have confirmed ligand-binding sites.^3–5^ To exploit these ligandable RNA pockets for novel activities, significant efforts have been dedicated to identifying RNA-binding small molecules using a variety of techniques ranging from rational design to high-throughput screening using binding and biological assays.^3,6^ Notably, contemporary efforts in the field have led to the discovery of small molecule ligands for a number of disease-relevant RNAs, forming the basis for a broader field of RNA-targeted drug discovery.^3,6^ Targeting RNAs with small molecules is still quite nascent, and the field would benefit greatly from development of new technologies that streamline elaboration of hits into leads for subsequent clinical translation.

Confirmation of target engagement inside cells is a crucial aspect of probe and drug discovery, for identification of both on- and off-target interactions.^7^ Over the past 10 years, several target engagement strategies have been reported for RNA-binding small molecules. Early efforts relied on the use of highly functionalized compounds bearing a nitrogen mustard alkylating agent and biotin for covalent crosslinking and affinity enrichment, respectively.^8–14^ More recently, photoaffinity-based crosslinking strategies have been employed for target engagement studies and transcriptome-wide profiling of RNA-small molecule interactions.^14–26^ Although these approaches have been used to successfully identify predicted target RNAs, none are able to elaborate in detail the sequence or structural details of small molecule binding site motifs. Being able to define the exact sequences engaged by small molecules is critical, as disease-relevant RNA transcripts tend to be very large (hundreds to thousands of nucleotides) and exhibit significant sequence heterogeneity, particularly in regulatory regions (e.g., 5′ and 3′ untranslated regions (UTR)).^27^ RNAs also exist within unique conformational ensembles inside and outside cells depending on RNA-binding proteins, RNA modifications, and other environmental differences.^1,28,29^ While binding site confirmation studies meant to validate on-target activity of a pharmacologically active small molecule ligand have largely been performed outside cells, the complex nature of RNA targets necessitates the need for early-stage validation inside cells to prioritize compounds for further optimization.

The use of RNA structure probing technologies, for example selective 2′-hydroxyl acylation analyzed by primer extension (SHAPE)-based approaches, have enabled experimental elaboration of RNA structural dynamics and provided tools for RNA-targeted drug discovery.^1,30–35^ SHAPE-MaP, which uses mutational profiling (MaP) as a sequencing readout of SHAPE reactivity, has a distinct advantage in that it allows for highly parallelized measurement of RNA flexibilities at single-nucleotide resolution both inside and outside cells.^32,33,36^ In MaP, chemical modifications (for example, SHAPE adducts) are “read through” by reverse transcriptase (RT) instead of triggering RT stops as in earlier SHAPE approaches. MaP read-through is enabled by altering the buffer conditions of select RT enzymes, which allows them to elongate past adducts by adding non-templated nucleotides to the cDNA product, resulting in a cDNA product with “mutations” at the site of chemical adducts. Mutation-containing cDNA can be sequenced at high depth to obtain mutation frequencies for every nucleotide in an amplified target, and mutational frequency is proportional to probe reactivity. To our knowledge, SHAPE-MaP has been rarely applied to the profiling of ligand-directed labeling of RNA targets in cells,^37^ and most target engagement studies have used RT stops to infer binding/labeling sites.^12,16,17,22,24,25^ Ideally, positions of RT stops can be used to infer reactive positions with nucleotide resolution, as in SHAPE; however, because RT stops can occur randomly, nucleotide-resolution assignment of reactivity remains a challenge. Using the preQ_1_ riboswitch as a model, herein we describe our chemistry solution for developing a probing strategy that can successfully be integrated with in-cell SHAPE-MaP.

## RESULTS AND DISCUSSION

### Design of an icSHAPE-based strategy for enabling target engagement analyses of cellular RNA-small molecule interactions

In RNA, only 2 chemical moieties distinguish it from DNA: the 2′-hydroxyl group (2′-OH) on the ribose ring and uridine as a nucleobase. The 2′-OH, present in every RNA nucleotide, has been exploited to create RNA-selective acylation chemistries due to its appropriate nucleophilicity for reaction with activated carbonyl compounds.^38,39^ This type of RNA biorthogonal chemistry has been adapted for many live-cell RNA applications,^39,40^ most notably, transcriptome-wide in-cell RNA structure probing through a technique known as *in vivo* click selective SHAPE, or icSHAPE.^31,41,42^ SHAPE reagents exhibit high reactivity at the 2′-OH of conformationally dynamic RNA regions, while low reactivity is observed at RNA nucleotides engaged in base-pairing or interactions with other macromolecules.^31,34,41^ Thus, SHAPE reactivities are most frequently used as additional restraints when modeling RNA secondary structure.^43^ By appending an azide to the SHAPE reagent, icSHAPE-reactive RNAs can be selectively enriched: the azide is labeled with a biotin-containing strained cyclooctene through a copper-free azide-alkyne “click” reaction, and labeled RNAs are subsequently captured by streptavidin-mediated affinity chromatography.^34,44^

While icSHAPE reagents (e.g., NAI-N_3_ and FAI-N_3_) can react with any RNA nucleotide if the 2′-OH is favorable to acylation, we became excited about the possibility that icSHAPE-based electrophiles could be elaborated as covalent probes for the validation of cellular RNA targets, as small molecules typically bind to RNAs at conformationally dynamic (flexible) sites.^45^ Moreover, progress in developing new RNA acylation probes has provided evidence that reactivity can be tuned through modification of the chemical structure of the acylating reagent.^46,47^ Indeed, concurrent to our studies, an acylation-based probing strategy termed “reactivity-based RNA profiling” (RBRP) was developed to identify RNA targets of approved drugs, and binding sites were mapped indirectly based on enrichment of RT stops.^48^ Also concurrently, cgSHAPE-seq (chemistry-guided SHAPE sequencing) was developed to identify the binding site of a small molecule on purified SARS-CoV-2 5′ UTR RNA, employing an acylation-based probing strategy and MaP.^49^ Our unique acylation chemistry takes a “best of both worlds” approach modifying an FAI-N_3_ icSHAPE reagent scaffold that retains the azide click chemistry handle (Figure 1A).^49^ FAI-N_3_ was selected as an icSHAPE warhead for further elaboration due to its reduced reactivity and longer half-life, which we predicted would enrich for true ligand-directed RNA labeling inside cells while enabling MaP-directed target identification (Figure 1B).^31,49–52^

**Figure 1.**
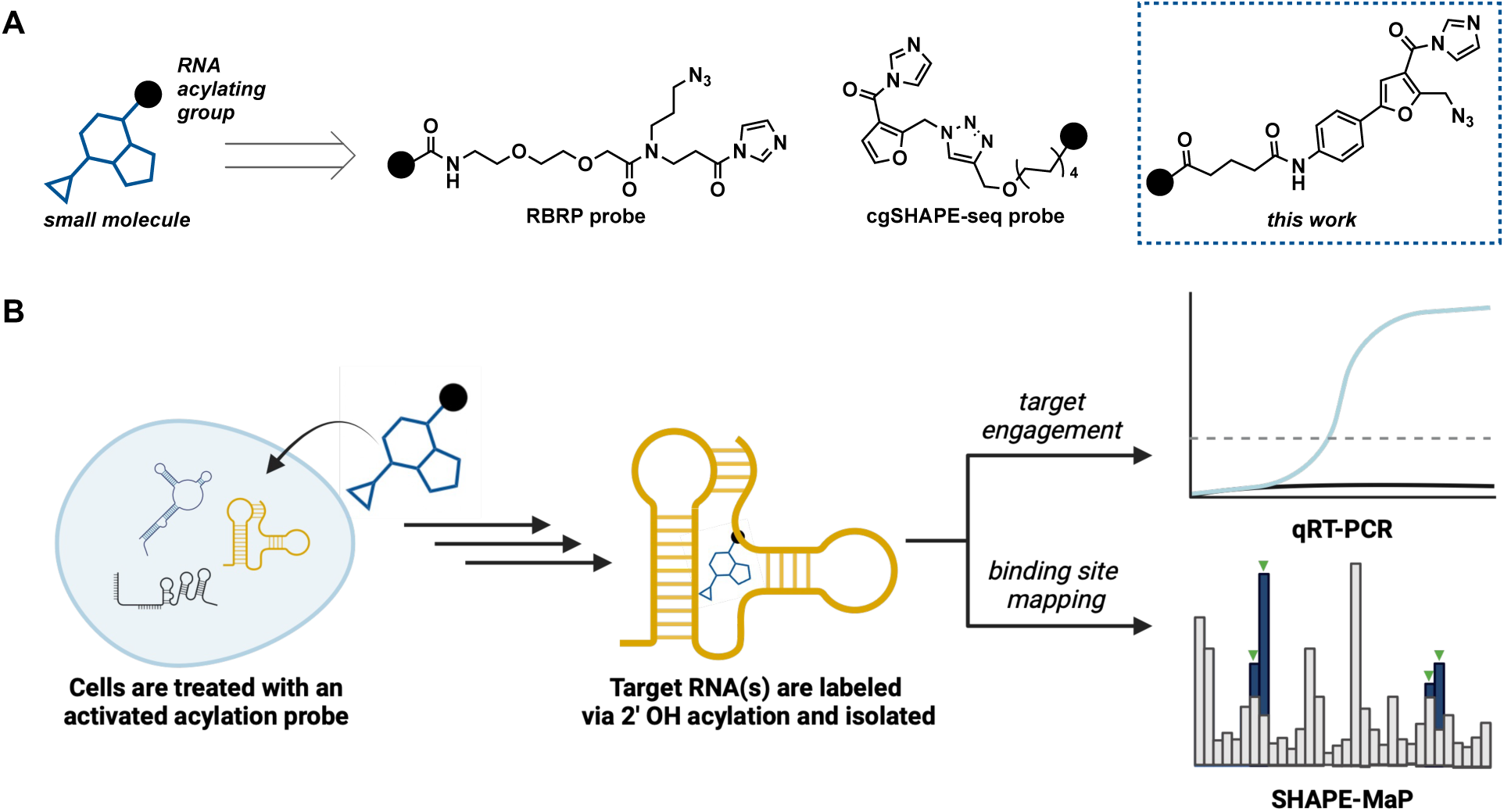
RNA acylation-based strategies for analyzing and discovering RNA-small molecule interactions. (A) Varying probe designs. (B) Workflow for using SHAPE-based acylation probes for in-cell investigation of target engagement and binding site mapping of RNA-small molecule interactions.

### Design and synthesis of a preQ_1_ icSHAPE-based probe

Riboswitches present an exceptional class of RNAs due to their highly structured natures and unique metabolite binding mechanisms.^53,54^ Riboswitch-small molecule interactions thus serve as excellent model systems based on their potent and selective nature, known bioactivity, and extensive previous investigation for biotechnology applications, including the development of photoaffinity-based profiling approaches.^20,21,26^ We selected the small molecule metabolite preQ_1_ as a model RNA-binding ligand (Scheme 1), which binds to preQ_1_ riboswitches to regulate gene expression in bacteria.^55^ The preQ_1_ riboswitch has been well characterized structurally, including using icSHAPE and *in vitro* SHAPE-MaP,^56,57^ and a photoaffinity-based preQ_1_ probe was used to discover cellular RNA targets from a mammalian cell line.^20^

**Scheme 1.**
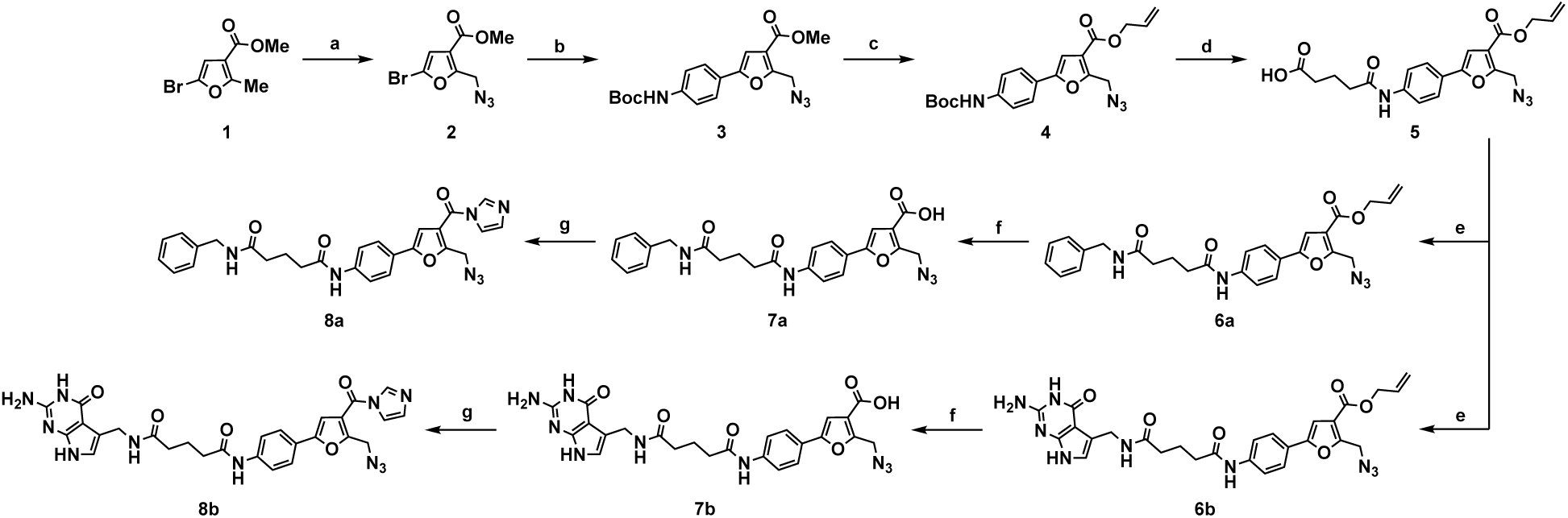
Synthesis of icSHAPE-based probes. Reagents and conditions: **a.** (i) NBS, AIBN, CCl_4_, 50 °C; (ii) NaN_3_, DMF, 23 °C, 83% over 2 steps; **b.** 4-(*N*-Boc-amino)phenylboronic acid pinacol ester, Pd(PPh_3_)_4_, K_2_CO_3_, 1,4-dioxane, 55 °C, 46%; **c.** (i) LiOH•H_2_O, 1,4-dioxane:H_2_O (3:1), 23 °C; (ii) allyl bromide, K_2_CO_3_, DMF, 0 °C to 23 °C, 55% over 2 steps; **d.** (i) TFA:CH_2_Cl_2_ (1:4), 0 °C; (ii) glutaric anhydride, DIPEA, DMAP, DMF, 23 °C; 82% over 2 steps; **e.** benzylamine or preQ_1_, DIPEA, HATU, DMF, 23 °C, 33% for **6b**; **f.** Pd(PPh_3_)_4_, pyrrolidine, DMF, 23 °C, 43% for **7a** (over steps **e.** and **f.)** and 68% for **7b**; **g.** CDI, DMSO, 23 °C, 76% for **8a** and 41% for **8b**.

To test our icSHAPE-inspired strategy, we synthesized preQ_1_ and control icSHAPE probes (Scheme 1). First, we prepared an FAI derivative containing an aryl amine handle for further elaboration (**4**). We envisioned that this aryl substitution could provide additional ν,ν-stacking interactions with an RNA substrate. The probe scaffold was subsequently modified to bear a glutaric acid linker (**5**) to enable differential display of the icSHAPE warhead and an RNA-binding ligand. **5** was then coupled to either benzylamine (**6a**) as a low-complexity control probe or preQ_1_ (**6b**). Ester deprotection and conversion to the activated acyl imidazole with 1,1’-carbonyldiimidazole (CDI) produced icSHAPE-based probes **8a** and **8b** in overall moderate yield.

### Live-cell confirmation of probe activity using a fluorescent reporter system

Prior to target engagement studies, we validated our preQ_1_ icSHAPE probe design by confirming its biological activity using a previously reported fluorescence reporter assay.^56–58^ The preQ_1_ riboswitch sequence from *Lactobacillus rhamnosus* (*Lrh*) was inserted into a plasmid expressing UV-excited green fluorescent protein (GFPuv) (Figure S1A). We positioned the riboswitch upstream of the GFP coding sequence such that it would be included in the 5′ UTR of expressed GFPuv RNA. The resulting *Lrh*-preQ_1_-GFPuv construct was then transformed into *ΔqueF E. coli* cells (Keio strain JW2765)^59^ that cannot biosynthesize the preQ_1_ metabolite. In the absence of a preQ_1_ riboswitch ligand, cells express GFPuv protein and produce fluorescence signal (Figure S1B); however, in the presence of a competent ligand, the ribosome binding site is sequestered in a riboswitch pseudoknot structure, preventing translation of GFPuv and consequently reducing fluorescence signal.^56^ PreQ_1_-GFPuv-transformed cells were grown in chemically defined media^58^ and treated with either probe, preQ_1_, or DMSO for 5 h at 37 °C before measuring fluorescence. We selected the non-activated allyl ester intermediate **6b** (Scheme 1) to test binding-based activity. The non-activated allyl ester does not react with RNA and instead reversibly binds to the riboswitch, ensuring that the probe will remain available throughout the multiple generations of bacterial turnover necessary for GFPuv protein production and detection. Importantly, from this analysis, we found that **6b** exhibited dose-dependent inhibition of fluorescence production when compared to DMSO-treated cells (Figure 2). The observed bioactivity of the preQ_1_-based probe was notably weaker than that of preQ_1_ (Figures 2 and S1; the reported EC_50_ for fold repression for preQ_1_ is 6.9 nM).^56^ However, reduced activity was not unexpected: the 7-aminomethyl group that we used for functionalization participates in electrostatic interactions with the riboswitch.^60,61^ Similar photoaffinity-based preQ_1_ probes using the same site of modification on the preQ_1_ scaffold also showed 50-fold weaker binding than preQ_1_ and further diminished potency in a transcription termination assay, equivalent to our observations.^20^ Based on the reproducible activity with orthogonal photoaffinity probes, we were confident we could confirm target engagement with our activated icSHAPE preQ_1_ probe.

**Figure 2.**
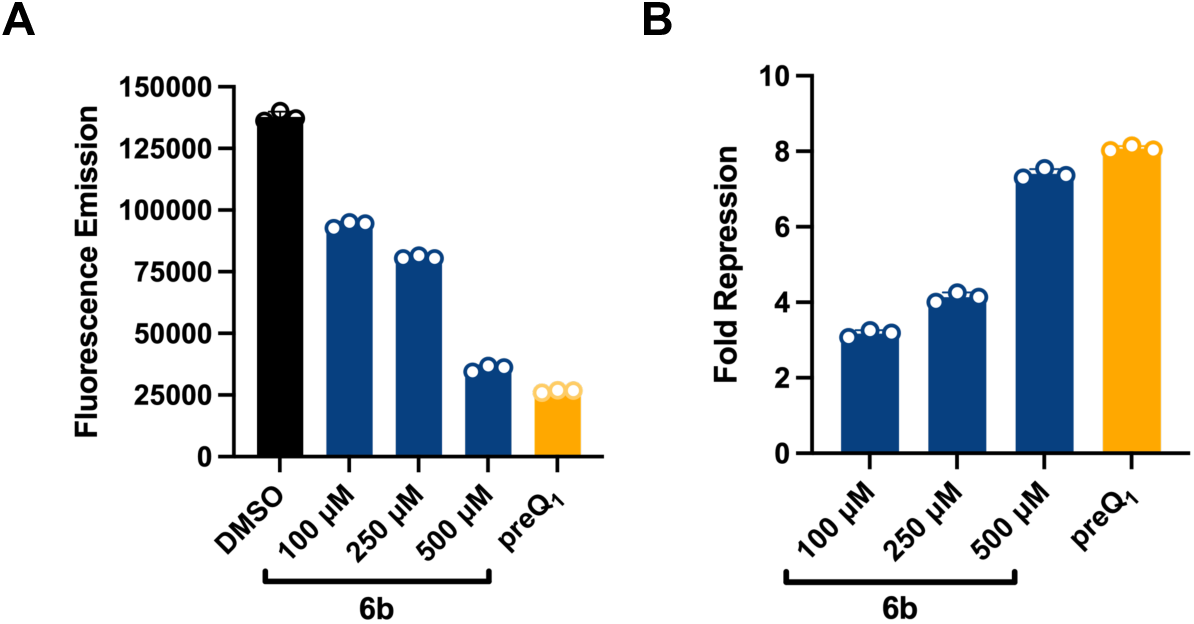
Bioactivity of non-activated preQ_1_ probe **6b** in a *Lrh*-preQ_1_-GFPuv reporter assay. (A) Fluorescence emission and (B) fold-repression following treatment of *Lrh*-preQ_1_-GFPuv-transformed JW2765 *E. coli* cells with **6b** or preQ_1_ (10 μM).

### Target engagement of the preQ_1_ riboswitch from live bacterial cells

Chemical crosslinking and isolation by pulldown (Chem-CLIP)^8,62^ strategies have confirmed target engagement between RNA targets-of-interest and small molecule probes.^9–11,13–16,18,20,21,23–26^ To determine if our icSHAPE-based probes were amenable to Chem-CLIP profiling in bacterial cells, we developed a pulldown pipeline for analysis of enriched RNA using qRT-PCR (Figure 3). *Lrh*-preQ_1_-GFPuv-transformed cells were grown for 1 h at 37 °C in the presence of acyl imidazole probes **8a** or **8b** (50 μM) or DMSO as a negative control. Total RNA was then isolated from lysed cells and subjected to copper-free click chemistry conditions via treatment with DBCO-PEG4-Biotin.^63^ Biotinylated RNAs from the mixture were enriched with streptavidin magnetic resin and eluted for analysis.

**Figure 3.**
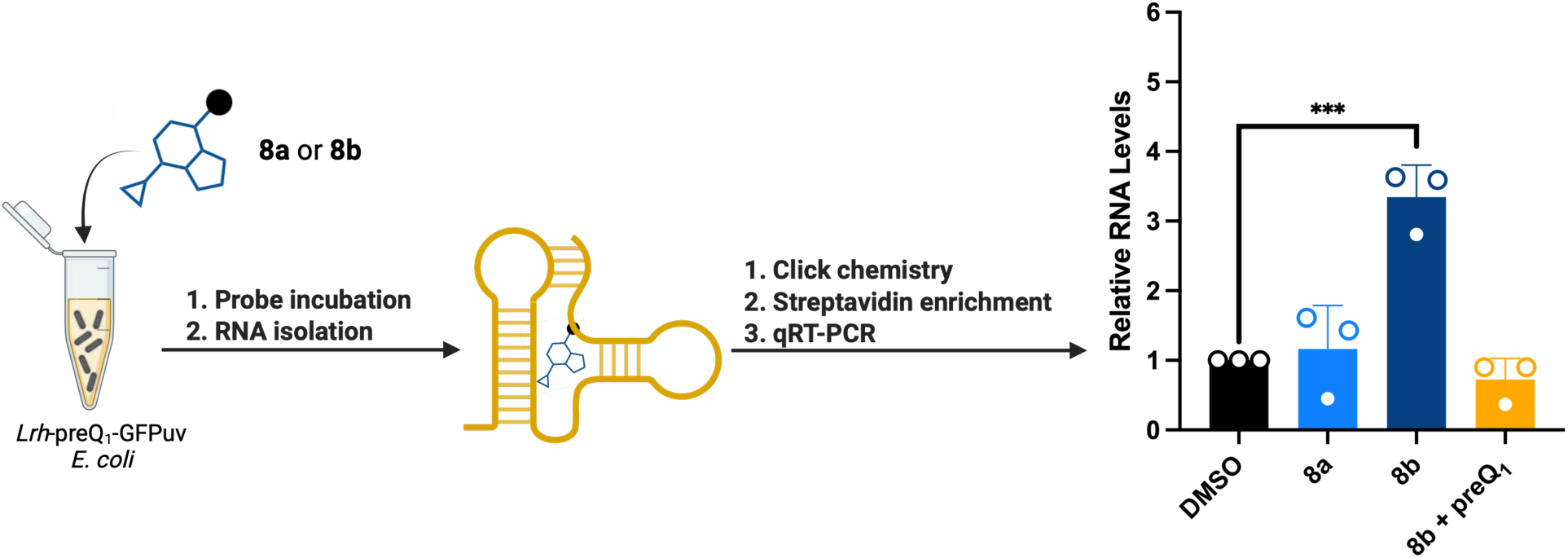
Chem-CLIP using icSHAPE-based probes for target engagement analysis using qRT-PCR. [**8a**] and [**8b**] = 50 μM. DMSO was used as a negative control. Competition studies were performed by co-treating with 10 μM preQ_1_. Statistical significance was determined using an unpaired, two-tailed Student’s *t* test. *** *p* = 0.0009.

For qRT-PCR analysis, cDNA was synthesized from streptavidin-enriched RNAs, and qPCR analysis was performed using gene-specific primers for *GFPuv*. While relative levels of GFPuv RNA were comparable from the streptavidin-enriched pool for both DMSO and low-complexity benzylamine control probe **8a**, excitingly, a 3.3-fold enrichment was observed with preQ_1_ icSHAPE probe **8b** (Figure 3). Notably, this level of enrichment is similar to that observed with photoaffinity-based riboswitch probes in live cell experiments.^21,26^ To demonstrate on-target enrichment, we performed a competitive analysis by co-treating cells with **8b** (50 μM) and preQ_1_ (10 μM). As expected for an on-target interaction, the presence of preQ_1_ abolished **8b**-dependent enrichment, suggesting competition for binding to the preQ_1_ riboswitch. Combined, these data demonstrate that an elaborated icSHAPE-based probe can retain cellular activity and target engagement, even with a highly structured RNA like the preQ_1_ riboswitch.

### Integration with in-cell SHAPE-MaP for binding site identification

As additional evidence of on-target activity, we explored the extent of binding and labeling of the preQ_1_ riboswitch in bacterial cells through mutational profiling (MaP). With MaP, specialized reverse transcription reads through modified nucleotides on RNA and deposits non-templated nucleotides in cDNA products at locations of adducts, which can be mapped by high throughput sequencing.^32,33^ Mutation rates observed during MaP are proportional to the level of a nucleotide’s reactivity to probe, and with our covalent probes, increased mutation rates would be indicative of sites of binding and labeling.

*Lrh*-preQ_1_-GFPuv-transformed cells were treated with icSHAPE preQ_1_ probe **8b** (50 μM) for 1 h at 37 °C and labeled RNAs were enriched using our streptavidin enrichment approach. We measured mutation rates from MaP RT of both probe-treated input and our streptavidin-enriched RNA, along with total RNA harvested from DMSO-treated cells, using primers specific to the 5′ end of the preQ_1_-GFPuv RNA reporter. While we treated with probe at concentrations multiple orders of magnitude below that of standard in-cell structure probing SHAPE-MaP experiments (50 μM of probe versus 10+ mM of a standard SHAPE reagent^44^), we observed a significant increase in observed mutation rates in the 5′ region of preQ_1_-GFPuv when comparing probe-treated cells to DMSO-treated cells (Figures 4A and S2). Further, MaP mutation rates from streptavidin-enriched RNA were significantly higher still (∼50%) than the probe-treated input (Figure S2), increasing confidence that our approach effectively enriches for probe-reactive RNA. The most reactive nucleotide (G82 in our Figures) lies in the pseudoknotted ribosome binding site, adjacent to a group of nucleotides in close three-dimensional proximity to the preQ_1_ binding site, based on atomic-resolution structure data (Figure 4A).^60^ G82 is about 20 Å from the binding site, a distance accommodated by the length of the probe linker. Nucleotides on the other end of the psuedoknot, A40 and U43, were also highly reactive (Figure 4A). Interestingly, we observed significant MaP signal in the downstream GFPuv coding sequence, especially C180 and C236 (Figure 4A). While C236 can be rationalized by its inclusion in a predicted base-paired stem close to the pseudoknot, it is difficult to predict how the C180 loop nucleotide might be positioned near the preQ_1_ binding site, although linker length and structural heterogeneity could enable a favorable conformation for reactivity.

**Figure 4.**
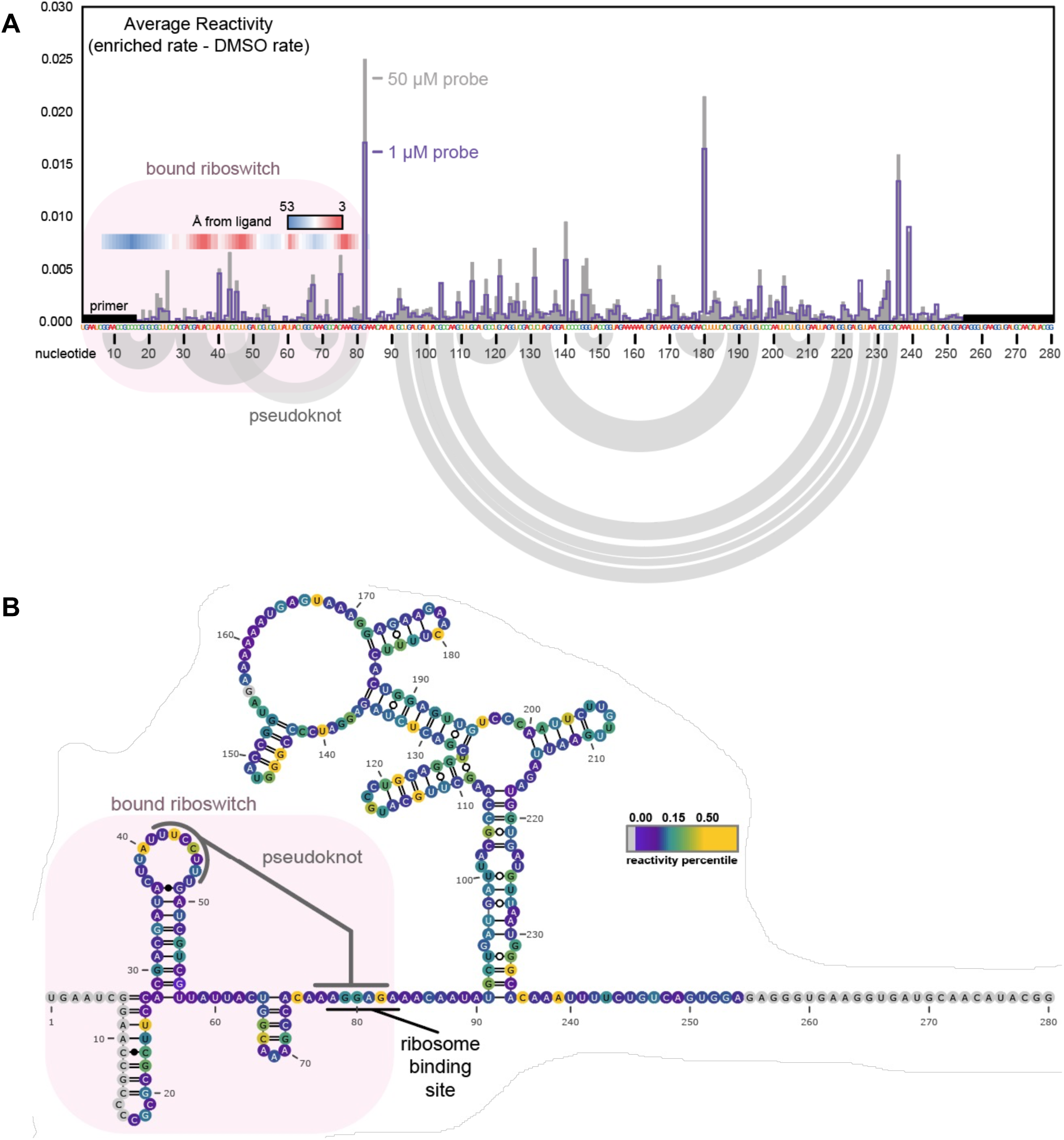
SHAPE-MaP analysis of icSHAPE preQ_1_ probe binding to the *Lrh*-preQ_1_ riboswitch in bacterial cells. (A) SHAPE mutation rates across the *Lrh*-preQ_1_-GFPuv transcript following treatment with 10 μM or 50 μM **8b**. (B) **8b** reactivities shown on the predicted structure of the *Lrh*-preQ_1_-GFPuv transcript.

We decided to vary the concentration of our probe even lower, in hopes of being able to unambiguously assign probe-binding nucleotides directly from MaP data. Even at 1 μM, MaP signal in the aptamer domain clearly defined regions responsible for engaging preQ_1_; however, significant reactivity was still observed in the nearby body of the GFP gene (Figure 4A). Plotting probe reactivity onto an RNA secondary structure model revealed that both paired and unpaired nucleotides were significantly reactive, suggesting that reactivity is likely more strongly related to binding proximity than to the intrinsic conformational flexibility of the unbound RNA (Figure 4B). That meaningful reactivity is even measurable at this concentration, approaching the K_d_ of the native ligand, speaks to the sensitivity of the designed probe.

To further confirm that labeling of the preQ_1_-GFPuv RNA was indeed proximity-directed and not a result of intrinsic reactivity of certain sequences to SHAPE reagents, we performed in-cell SHAPE-MaP using FAI-N_3_^50^. We measured SHAPE reactivity of the riboswitch in cells treated with preQ_1_ or allyl ester-containing preQ_1_ probe **6b**. Notably, we observed highly similar SHAPE reactivity patterns of mutations between the samples (Figures 5 and S3) and with previously reported SHAPE-based analyses of preQ_1_ bound to the *Lrh*-preQ_1_ riboswitch.^56,57^ These results provide greater confidence that **6b** binds to the expected metabolite binding pocket: no significant deviations within either the riboswitch aptamer or the GFPuv coding sequence were observed between the samples, indicating that the RNA folds similarly in the presence of either ligand. Importantly, MaP signal from **8b** treated and enriched RNA was slightly anti-correlated with FAI SHAPE-MaP signal from **6b**-bound RNA (Figure S3), supporting that **8b** reactive sites are more likely probe proximity-induced rather than highly reactive with FAI-N_3_. In fact, all nucleotides highly reactive to **8b** were ultimately unreactive to FAI-N_3_, whether bound by preQ_1_ or **6b** (Figure 5). As the linker incorporated into the **8b** probe is flexible and is displayed outside of the preQ_1_ binding pocket (Figure S4), we hypothesize that this contributes to the diversity of mutation and labeling sites. Future studies will investigate the impact that linker chemistry has on probe labeling reactivity and specificity.

**Figure 5.**
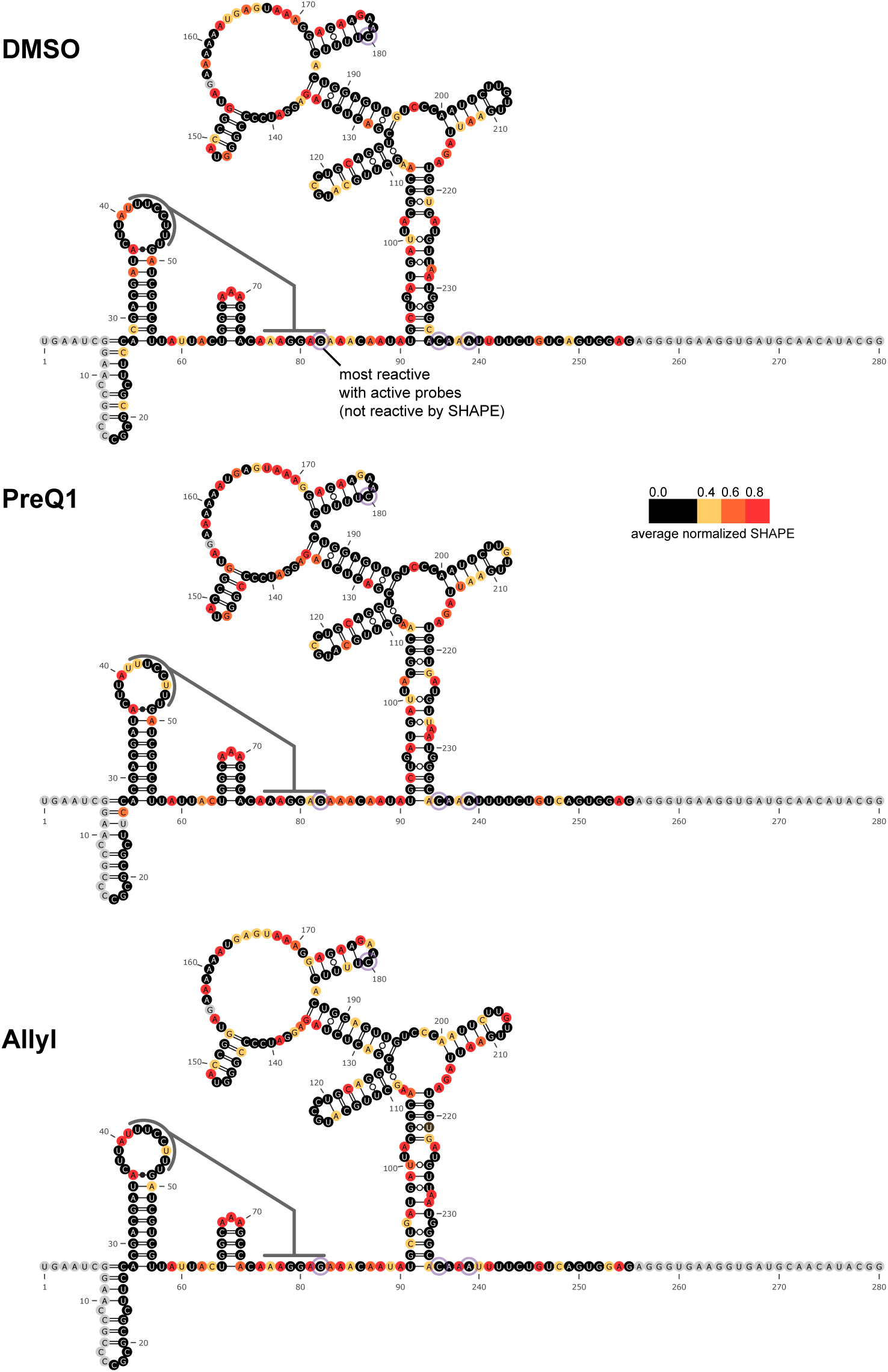
Average normalized FAI-N_3_ SHAPE reactivity from three replicates across the *Lrh*-preQ_1_-GFPuv transcript 5′ end following in-cell treatment with DMSO (top), preQ_1_ (5 μM) (middle), or preQ_1_ probe **6b** (50 μM) (bottom). **8b**-reactive sites are circled.

## CONCLUSION

The ability to manipulate and investigate cellular RNA-small molecule interactions has broad significance in biotechnological applications and human health. As the field continues to grow, new tools are needed to support these efforts. Inspired by established RNA acylation chemistry, we developed a chemical probing approach that enabled interrogation of the interaction between preQ_1_ and the preQ_1_ riboswitch at near nucleotide resolution in live bacterial cells. Notably, the ability to integrate chemical profiling with SHAPE-MaP as highlighted herein has been a challenge in the field with analogous photoaffinity-based probes.^17^ Based on our promising results, we anticipate broader adoption of this methodology for RNA target discovery and the validation of RNA-small molecule interactions in cells. Further advancement of this technology for this type of profiling will be reported in due course.

## ASSOCIATED CONTENT

### Supporting Information

The Supporting Information is available free of charge at… Methods, supplemental figures, NMR spectra (PDF)

## AUTHOR INFORMATION

Authors

**José A. Reyes Franceschi** – Program in Chemical Biology, University of Michigan, Ann Arbor, Michigan 48109, USA

**Emilio L. Cárdenas** - Department of Medicinal Chemistry, College of Pharmacy, University of Michigan, Ann Arbor, Michigan 48109, USA

**Brandon J. C. Klein** - Department of Medicinal Chemistry, College of Pharmacy, University of Michigan, Ann Arbor, Michigan 48109, USA

Notes

L. G. is an advisor to and holds equity in Skyhawk Therapeutics and Atomic AI.

## ACKNOWLEDGMENTS

This work was supported by the NIH (R01 GM132342 and R35 GM153185 to A.L.G., R01 GM155542 to C.A.W.), the Dr. Ralph and Marian Falk Medical Research Trust (A.L.G.), the University of Michigan College of Pharmacy Upjohn Award (A.L.G.), and a Rackham Merit Fellowship from the University of Michigan Rackham Graduate School (J.R.F.). Figures 1b, 3, S1A, and the table of contents graphic were created using Biorender.

## ABBREVIATIONS

SHAPE: selective 2′-hydroxyl acylation analyzed by primer extension
SHAPE-MaP,: SHAPE with mutational profiling
icSHAPE: *in vivo* click selective SHAPE
UTR: untranslated region
RT: reverse transcriptase

## TABLE OF CONTENTS GRAPHIC

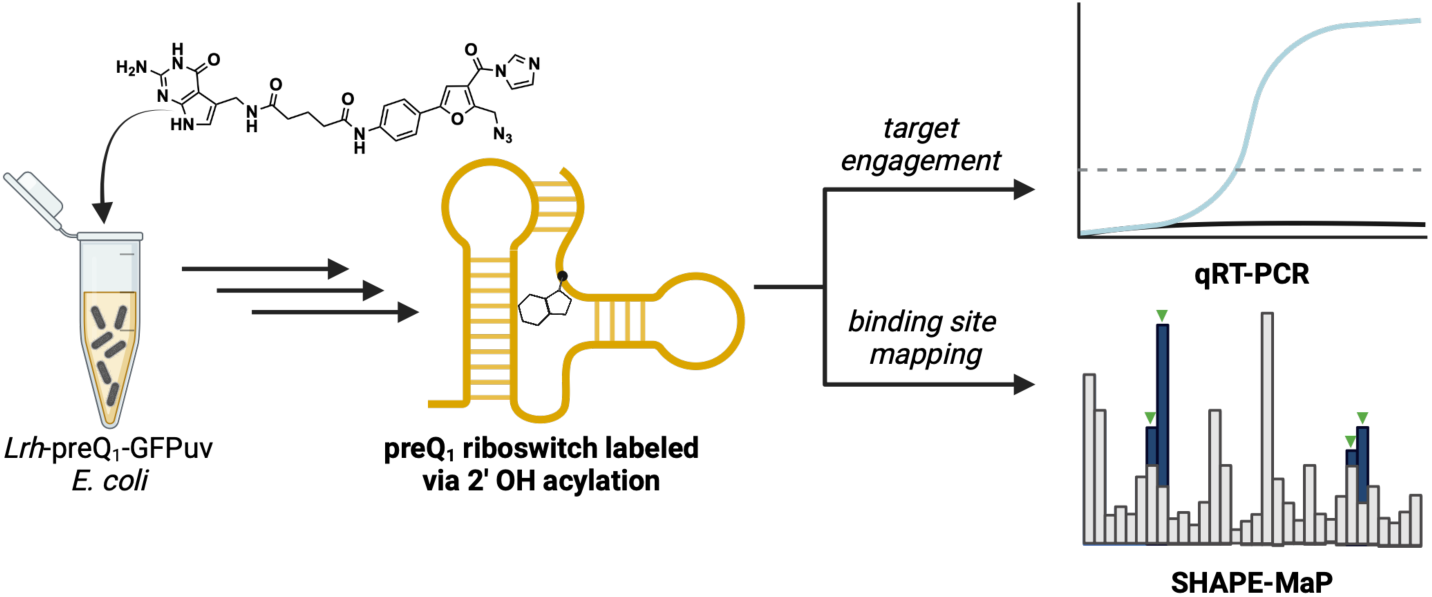

