## Supplemental Information for "SHAPE-based chemical probes for studying preQ_1_-RNA interactions in living bacteria"

**A. Biology Methods** Pages S2–S6

**B. Supplementary Figures** Pages S7–S8

**C. Synthetic Methods** Pages S9–S14

**D. NMR Spectra** Pages S15–S24

**E. References** Page S25

#### B. Biology Methods

**Construction of a GFPuv *Lrh* preQ$_1$-II Riboswitch reporter plasmid.** Plasmid construction followed that previously described.[1] In brief, a synthetic DNA insert comprising the *Lactobacillus rhamnosus* (*Lrh*)-preQ$_1$ riboswitch sequence (5'-AAGGCTCGTATAATAACTAT TAACTGAATCGGAACCGCCCCGCGCGCTTCCACGACGATACTTATTTCCTTTGATCG TCGTTATTACTGGCAAAGCCACAAAGGAGAAACAATATGCTGATGATTACGCC-3') was prepared in plasmid pUC57 (GenScript) and subcloned into the pEnv8 (GAAA) vector (AddGene) using NsiI-HF and HindIII-HF (NEB) restriction sites. The resulting plasmid, *Lrh*-preQ$_1$-GFPuv, was amplified in DH5α *E.coli* cells in Luria-Bertani broth (LB) supplemented with 100 µg/mL ampicillin.

**Fluorescence reporter assay.** The *Lrh*-preQ$_1$-GFPuv construct was transformed into competent *E. coli* strain JW2765 (Horizon Discovery)[2] which is *ΔqueF* and does not produce preQ$_1$ as previously described.[1] Cells were plated and grown on agar plates containing chemically-defined CSB media (0.033 M $NaH_2PO_4$, 0.086 M $K_2HPO_4$, 0.015 M $(NH_4)_2SO_4$, 0.005 M sodium citrate pH 6.0, 0.492 mM $MgSO_4$, 0.109 mM $FeCl_3$, 0.0283 mM $ZnCl_2$, 0.0248 mM $CoCl_2$, 0.0138 mM $Na_2MoO_4$, 0.0201 mM $CaCl_2$, 0.0219 mM $CuCl_2$, 0.0234 mM $MnCl_2$, 0.0239 mM $H_3BO_3$, and 6% (*w/v*) D-glucose)[3] supplemented with 100 µg/mL ampicillin (CSB-*amp*) overnight at 37 °C.

For the assay, colonies were selected and grown in 5 mL of liquid CSB media at 37 °C overnight. Cells were diluted to an $OD_{600}$ of 0.05 in fresh CSB-*amp* supplemented with either preQ$_1$ (10 µM) or **6b** (50−500 µM) at a final DMSO concentration of 5% and grown for ~5 h to mid-log phase ($OD_{600}$ ~0.6). 100 µL of each bacteria treatment was then plated in a black, clear-bottomed 96-

well plate and the $OD_{600}$ was measured for normalization. GFPuv fluorescence was then measured using a BioTek Cytation 3 plate reader ($\lambda_{excitation}$ = 395 nm; $\lambda_{emission}$ = 510 nm). Raw fluorescence readings were normalized using $OD_{600}$ measurements and background corrected; the background was calculated by obtaining fluorescence readings of non-transformed JW2765 cells, experiments were carried out in biological triplicates. The fold emission repression was calculated using the following formula: $y(x)=[1-preQ1(x)/ preQ1(0)]\cdot 10$, where x is the concentration of $preQ_1$ used.[1]

**Isolation of probe-enriched RNAs from live bacterial cells.** *Lrh*-$preQ_1$-GFPuv-expressing JW2765 cells were grown in 5-mL cultures overnight with CSB-*amp* media. Bacteria were pelleted at 3,900 rpm at 4 °C for 15 min, the media was aspirated, and the pellets were resuspended in 5-mL of fresh CSB media. The cell suspension was then divided into 1-mL samples, and each sample was supplemented with 50 µM final concentration of **8a** or **8b** (0.1% DMSO) or 0.1% DMSO as a negative control. After the addition of the new media, the cells were incubated for 1 h at 37 °C. Finally, RNAs were extracted from lysate using RNAzol® RT (Sigma). Total RNA extraction was performed following the manufacturer's protocol. Of note, RNA samples can be stored at -80 °C until needed.

Total RNA was concentrated to 100 µL and DNAse-treated using the Zymo RNA Clean & Concentrator™-5 kit following the manufacturer's instructions. Prior to enrichment, 1 µL of each sample (minimum 100 ng total RNA) was set aside to be used as an input control for future sequencing or qRT-PCR experiments. Concentrated samples then subjected to click chemistry. RNAs were diluted in 1× PBS pH 7.4 (125 µL) and treated with DBCO-PEG4-Biotin (Vector Laboratories) (25 µL of 5 mM, 10% DMSO). The reaction mixture was incubated at 37 °C for 2 h and purified using the Zymo RNA Clean & Concentrator™-5 kit.[4] RNAs were eluted in 2× 50 µL of RNAse free water. Eluted RNAs were then subjected to affinity enrichment using streptavidin resin (Dynabeads™ MyOne™ Streptavidin C1). Samples were diluted to 400 µL in bead binding buffer (50 mM citrate, pH 6.0, 1 mM EDTA) and incubated with streptavidin beads (100 µL of beads prewashed 3× with binding buffer) for 1 h at room temperature while spinning on a rotation rack. After incubation, beads were washed 2× with 500 µL of wash buffer (10 mM citrate, pH 6.0, 1 mM EDTA, 4 M NaCl, 0.25% Tween), 2× with 500 µL of 1× PBS (Invitrogen), and 2× with 500 µL of 70 °C water.[4] Washed beads were eluted 2× using 50 µL of a 70 °C 6M guanidine-HCl solution for 10 min. Combined elutions were purified using the Zymo RNA Clean & Concentrator™-5 kit. The concentration of eluted RNAs was measured using the Quant-it™ RiboGreen RNA quantification assay (Invitrogen). Samples were stored at -80 °C until needed for downstream analysis.

**qRT-PCR analysis of isolated RNAs.** Isolated RNAs were reverse transcribed into cDNA using the RevertAid First Strand cDNA Synthesis Kit following the manufacturer's instructions. qPCR analysis was performed on a QuantStudio™ 5 thermocycler (Thermo Scientific) in 384-well plate format using the PowerUp™ SYBR™ Green Master Mix for qPCR and gene specific forward and reverse primers for GFPuv. Fold enrichment was calculated using Sigma Aldrich method for Relative Quantification for Chip-qPCR.[5]

| **Primer** | **Sequence (5′…3′)** |
|---|---|
| GFPuv Reverse | CCATCTAATTCAACAAGAATTGGGACAAC |

| | |
|---|---|
| GFPuv Forward | GGTCCTTCTTGAGTTTGTAAC |

**In-cell acylation for total cell SHAPE-MaP.** *Lrh*-pre$Q_1$-GFPuv-expressing JW2765 cells were grown in 5-mL cultures overnight with CSB-*amp* media. Bacteria were pelleted at 3,900 rpm at 4 °C for 15 min, the media was aspirated, and the pellets were resuspended in 5 mL of fresh CSB media. The cell suspension was then divided into 1-mL samples, and each sample was supplemented with 5 μM final concentration of pre$Q_1$, 50 μM final concentration of **6b** (0.1% DMSO), or 0.1% DMSO as a negative control. After the addition of the new media, the cells were incubated for 1 h at 37 °C. Following incubation, cells were pelleted at 3,900 rpm at 4 °C for 15 min and resuspended in 1× PBS pH 7.4. Cells were then treated with either 100 mM final concentration of FAI-$N_3$ (10% DMSO) or 10% DMSO and incubated at 37 °C for 15 min. Reactions were quenched with 100 mM final concentration DTT, and cells were pelleted one final time at 3,900 rpm at 4 °C for 15 min. RNAs were extracted from lysates using RNAzol® RT (Sigma). Total RNA extraction was performed following the manufacturer's protocol. Total RNA was concentrated to 100 μL and DNAse-treated using the Zymo RNA Clean & Concentrator™-5 kit following the manufacturer's instructions. RNA aliquots were stored in -80°C until needed for MaP cDNA library prep.

**SHAPE-MaP.** Isolated RNAs from treatment with **8b** and a DMSO input RNA or total RNA extracted from treatments with **6b**, pre$Q_1$, or DMSO, were subjected to SHAPE-MaP experiments in biological triplicate. MaP-RT of the pre$Q_1$ riboswitch of the *Lrh*-pre$Q_1$-GFPuv construct was performed in 10 μL total volume. 5 μL of eluted enriched RNA was denatured at 70 °C for 5 min in a mixture containing 20 nmol of each dNTP base and an RT primer complementary to the 3′

end of the RNA. After 5 min, the mixture was immediately placed at 4 °C for 2 min. 9 µL of freshly made 2.22× MaP buffer (111 mM Tris-HCl [pH 8.0], 167 mM KCl, 13.3 mM $MnCl_2$, 22 mM DTT, and 2.22 M betaine) was added to the mixture, which was then equilibrated at 25°C for 2 min. Subsequently, 1 µL (200 units) of SSIIRT (Invitrogen) was added to the reaction, which was then placed in a thermocycler with the following settings: 10 min at 25 °C, 90 min at 42 °C, 10 cycles of 2 min at 42 °C and 2 min at 50 °C, and finally 10 min at 70 °C to heat inactivate the SSIIRT. $preQ_1$ specific forward and reverse primers were designed and used to perform multiplex PCRs following the manufacturers specifications for Q5 Hot-start Master mix. We then performed secondary PCR to add i5/i7 barcodes needed for multiplexing. Libraries were sequenced as paired-end 2 × 250 read multiplex runs on a MiSeq instrument. SHAPE reactivities were derived using the ShapeMapper2 pipeline.[6]

| **Primer** | **Sequence (5′…3′)** |
|---|---|
| $preQ_1$ RT | CCGTATGTTGCATCACCTTC |
| $preQ_1$ Reverse | TCGTCGGCAGCGTCAGATGTGTATAAGAG-ACAGNNNNNCCGTATGTTGCATCACCTTC |
| $preQ_1$ RT Forward | GTCTCGTGGGCTCGGAGATGTGTATAAG-AGACAGNNNNNTGAATCGGAACCGCCC |

**Data and Statistical Analysis.** All data was analyzed using GraphPad Prism version 9.5.1 for Mac OS (GraphPad Software, www.graphpad.com). Two-sided t tests were performed using Prism; equal variance between samples being compared was established.

#### B. Supplementary Figures

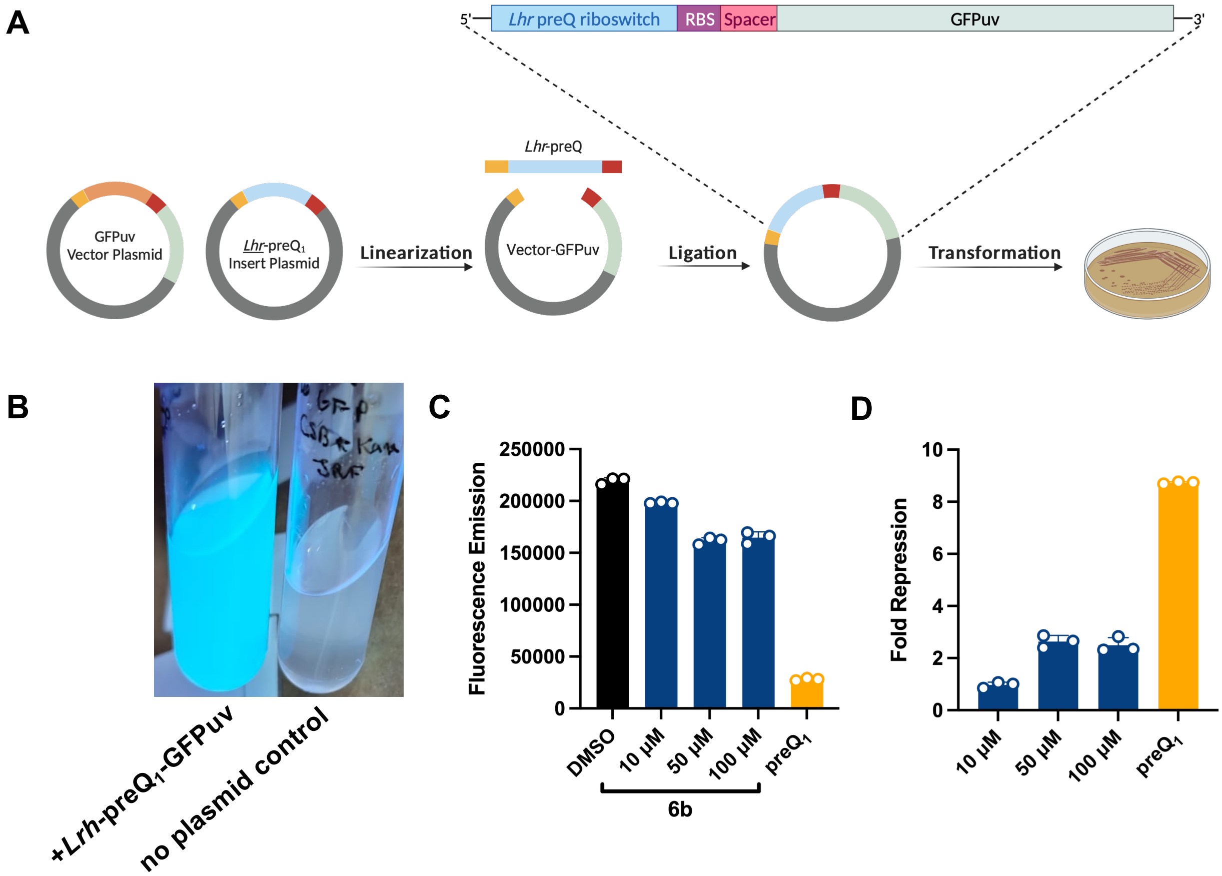


**Figure S1.** Reporter assay to characterize preQ$_1$ riboswitch activity. (A) Construction of the *Lrh*-preQ1-GFPuv plasmid. (B) Detection of GFPuv fluorescence in transformed JW2765 *E. coli* cells using a hand-held UV lamp (365 nm). (C) Fluorescence emission following treatment of *Lrh*-preQ1-GFPuv-transformed JW2765 *E. coli* cells with **6b** or preQ$_1$ (10 µM). (D) Fold fluorescence repression following treatment of *Lrh*-preQ1-GFPuv-transformed JW2765 *E. coli* cells with **6b** or preQ$_1$ (10 µM).

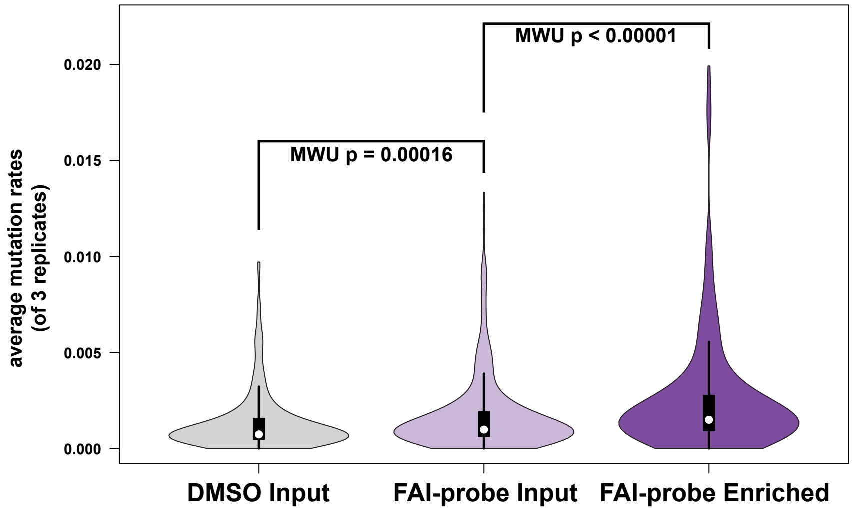


| | DMSO Input | FAI Input | FAI Enrichment |
|---|---|---|---|
| Upper whisker | 0.003096 | 0.003825 | 0.005437 |
| 3rd quartile | 0.001576 | 0.001927 | 0.00279933 |
| Median | 0.00073067 | 0.000982 | 0.00149217 |
| 1st quartile | 0.00046867 | 0.000605 | 0.00091667 |
| Lower whisker | 0 | 0 | 0 |
| Nr. of data points | 238 | 238 | 238 |

**Figure S2.** Violin plot summarizing mutation rates of the 5′ region of preQ1-GFPuv from indicated treatments in transformed *E. coli* cells. Box plot values are displayed. P-values from one-sided Mann-Whitney U-tests (MWU) are indicated.

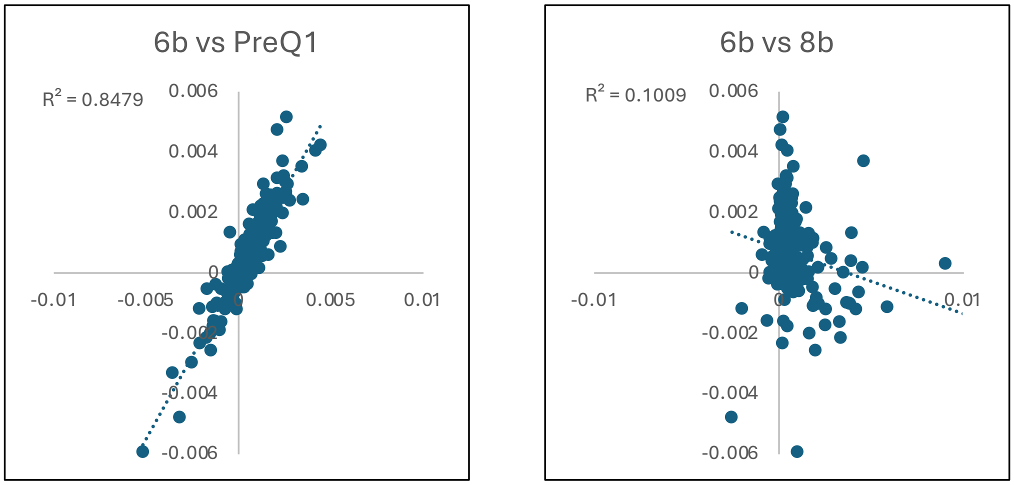


| | | |
|---|---|---|
| PEARSON coefficient | 0.920799 | -0.31766 |
| Comparison | PreQ1 : 6b | 6b : 8b |

**Figure S3.** Plots of correlation between indicated dataset mutation rates. Pearson's *r* and $R^2$ are labeled.

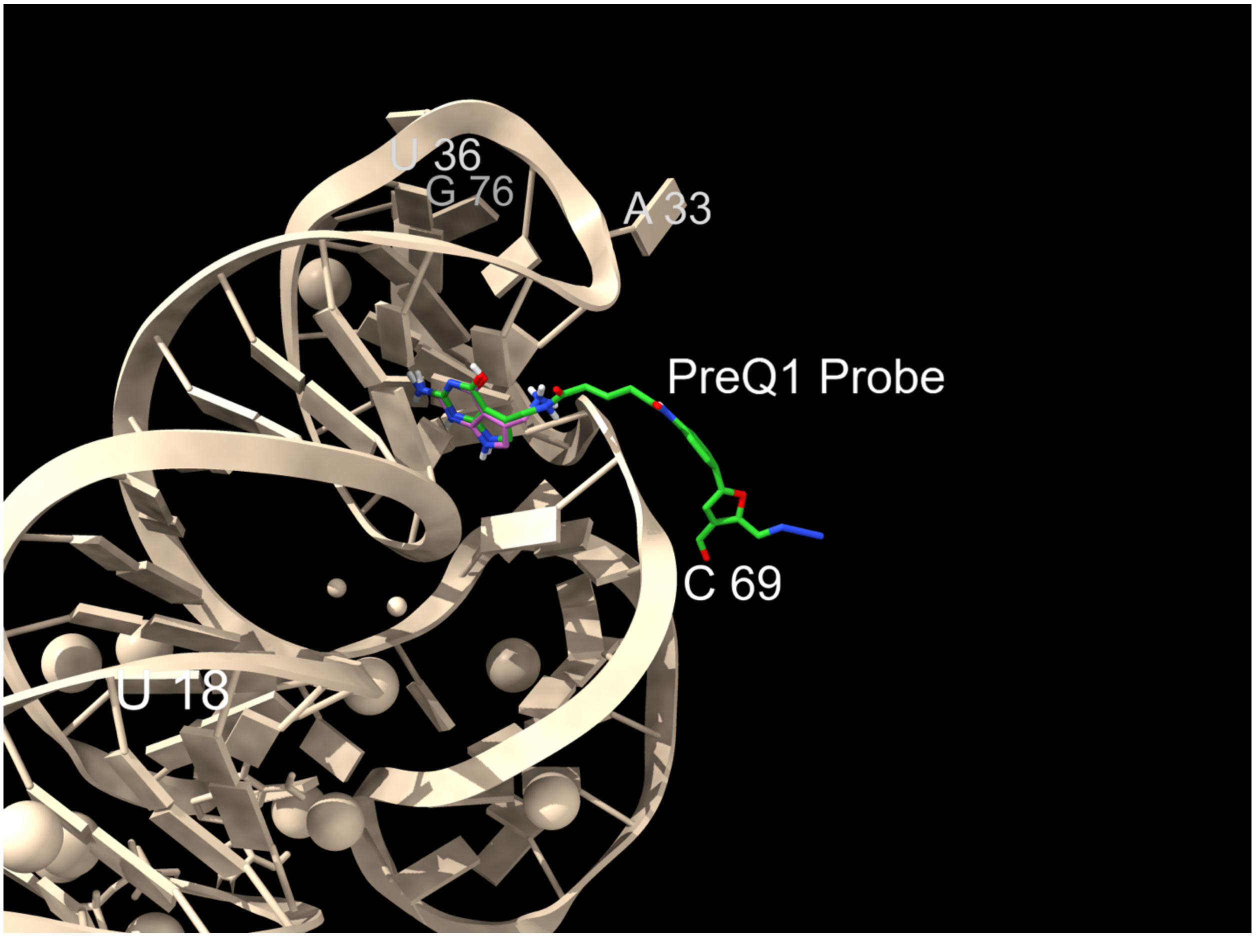


**Figure S4.** Overlay of preQ$_1$ probe on the crystal structure of preQ$_1$ bound to the *Lrh*-preQ$_1$ riboswitch (PDB 4JF2). PreQ$_1$ is shown in purple. PreQ$_1$ probe in shown in green. Highly reactive sites with probe **8b** are labeled (Figure 4).

#### C. Synthesis Methods

**General Chemistry Materials and Methods.** All purchased solvents and reagents were used without further purification. Reactions were monitored by thin-layer chromatography (TLC) carried out on 0.25-mm SiliCycle silica gel plates (60F-254) using UV-light (254 nm). Flash chromatography was performed using SiliaFlash P60 silica gel. Analytical RP-HPLC was performed using an Agilent 1260 Infinity HPLC equipped with an ZORBAX Eclipse XDB-C18 colun (4.6 x 150 mm; 5 μm) with detection at 254 nm. Semi-preparative HPLC was carried out on Agilent 1260 Infinity HPLC equipped with a PrepHT XDB-C18 column (21.2 x 150 mm; 5 μm) with detection at 254 nm). General Method A: 10–45% acetonitrile/$H_2O$ (0.1% FA) over 5 min at 12 mL/min. General Method B: 5–95% acetonitrile/$H_2O$ over 30 min at 12 mL/min. NMR spectra were obtained using 300 MHz Bruker and 400 MHz Bruker instruments calibrated using a solvent peak as an internal reference. Chemical shifts (*δ* values) are reported in parts per million and are referenced to the deuterated residual solvent peak. Mass spectrometry (MS) data was obtained using an Agilent 6230 TOF LC/MS spectrometer using ESI ionization with an accuracy of 2 ppm. All compounds were found to be >95% pure by HPLC analysis.

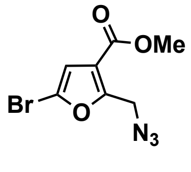


**2**

*Methyl 5-bromo-2-(azidomethyl)furan-3-carboxylate* (**2**). Methyl 5-bromo-2-methylfuran-3-carboxylate (**1**) (2.0 g, 9.17 mmol) was diluted in $CCl_4$ (0.2 M, 45 mL), treated with NBS (1.8 g, 10.1 mmol) and AIBN (76 mg, 0.46 mmol), and warmed to 50 ºC. After 16 h, the reaction mixture was cooled to 23 ºC, diluted with $CH_2Cl_2$ (200 mL), and washed with $NaHCO_{3(sat)}$ (100 mL), $H_2O$ (100 mL), and brine (100 mL). The organic layer was subsequently isolated, dried over $Na_2SO_4$, and concentrated to dryness to provide the crude product which was used without further purification. The crude analogue was diluted in DMF (0.3 M, 30 mL) and treated with $NaN_3$ (18.2 mmol, 1.18 g) at 23 ºC. After 24 h, the reaction mixture was concentrated, diluted with EtOAc (200 mL), and washed with $H_2O$ (100 mL) and brine (100 mL). The organic layer was dried over $Na_2SO_4$ and concentrated to dryness, and the isolated crude product was purified by column chromatography (3% EtOAc/Hexanes) to provide **2** (1.9 g, 83% over 2 steps). $^1$H NMR (400 MHz, $CDCl_3$) δ 6.68 (s, 1H), 4.63 (s, 2H), 3.88 (s, 3H); $^{13}$C NMR (101 MHz, $CDCl_3$) δ 155.95, 153.85, 123.32, 118.79, 112.51, 51.96, 45.26.

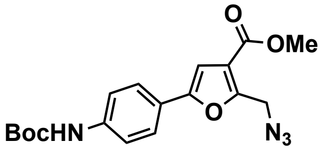

**3**

*Methyl 2-(azidomethyl)-5-(4-((tert-butoxycarbonyl)amino)phenyl) furan-3-carboxylate* (**3**). **2** (730 mg, 2.84 mmol) was diluted in 1,4-dioxane (0.06 M, 47 mL) and treated with 4-(*N*-Boc-amino)phenylboronic acid pinacol ester (1.0 g, 3.13 mmol) and $Pd(PPh_3)_4$ (164 mg, 0.14 mmol) at 23 ºC. After 5 min, a solution of $K_2CO_{3(aq)}$ (0.2 M, 28 mL) was added to the reaction mixture which was subsequently warmed to 55 ºC. After 5 h, the reaction was cooled to 23 ºC, passed through a pad of celite, and washed with EtOAc (100 mL). The filtrate was concentrated, diluted further with EtOAc (200 mL), and washed with $H_2O$ (100 mL) and brine (100 mL). The isolated organic layer was dried over $Na_2SO_4$, concentrated to dryness, and purified by column chromatography (10–15% EtOAc/Hexanes) to provide **3** (460 mg, 46%). $^1$H NMR (400 MHz, $CDCl_3$) δ 7.60 (d, $J$ = 7.3 Hz, 2H), 7.44 (d, $J$ = 8.3 Hz, 2H), 4.70 (s, 2H), 3.89 (s, 3H), 1.54 (s, 9H); $^{13}$C NMR (101 MHz, $CDCl_3$) δ 163.44, 154.05, 153.13, 152.55, 138.76, 124.96, 124.12, 118.55, 118.13, 104.42, 80.82, 51.82, 45.71, 28.31.

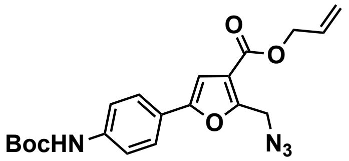

**4**

*Allyl 2-(azidomethyl)-5-(4-((tert-butoxycarbonyl)amino)phenyl) furan-3-carboxylate* (**4**). **3** (630 mg, 1.69 mmol) was diluted in a solution of 1,4-dioxane:$H_2O$ (3:1, 0.15 M, 12 mL) and treated with $LiOH•H_2O$ (142 mg, 3.38 mmol) at 23 ºC. After 17 h, the reaction mixture was concentrated to remove the organic volatiles, diluted further with $H_2O$ (100 mL), and acidified with 1N $HCl_{(aq)}$ (pH ~1–2). The aqueous layer was extracted with a solution of 5% $MeOH/CH_2Cl_2$ (3×200 mL), and the organic layer was dried over $Na_2SO_4$ and concentrated to dryness to provide the crude product that was used without further purification. The crude furoic acid was diluted in DMF (0.05 M, 30 mL) and treated with $K_2CO_3$ (583 mg, 4.22 mmol) and allyl bromide (292 µL, 3.38 mmol) at 0 ºC. After 15 minutes, the reaction was removed from the ice-water bath and allowed to reach 23 ºC. After 16 h, the reaction mixture was concentrated, diluted in EtOAc (200 mL), and washed with $NaHCO_{3(sat)}$ (100 mL), $H_2O$ (100 mL) and brine (100 mL). The isolated organic layer was dried over $Na_2SO_4$, concentrated to dryness, and purified by column chromatography (10% EtOAc/Hexanes) to provide **4** (370 mg, 55% over 2 steps). $^1$H NMR (400 MHz, $CDCl_3$) δ 7.63 (d, $J$ = 9.1 Hz, 2H), 7.44 (d, $J$ = 8.6 Hz, 2H), 6.89 (s, 1H), 6.55 (bs, 1H), 6.04 (m, 1H), 5.42 (dq, $J$ = 17.2, 1.6 Hz, 1H), 5.38 (dq, $J$ = 10.3, 1.3 Hz, 1H), 4.81 (dt, $J$ = 6.0, 1.4 Hz, 2H), 4.72 (s, 2H), 1.56 (s, 9H); $^{13}$C NMR (101 MHz, $CDCl_3$) δ 162.66, 154.07, 153.26, 152.53, 138.74, 131.87, 124.99, 124.14, 118.80, 118.54, 118.16, 104.43, 80.86, 65.45, 45.69, 28.32.

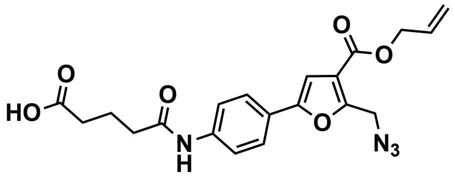

**5**

*5-((4-(4-((Allyloxy)carbonyl)-5-(azidomethyl)furan-2-yl) phenyl)amino)-5-oxopentanoic acid* (**5**). **4** (350 mg, 0.87 mmol) was diluted in a mixture of TFA:$CH_2Cl_2$ (1:4, 0.05 M, 15 mL) at 0 ºC. After 1 h, the reaction mixture was concentrated to dryness to provide the crude product which was used without further purification. The crude allyl furan carboxylate was subsequently diluted in DMF (0.1 M, 9 mL), and treated with DIPEA (768 µL, 4.3 mmol), DMAP (11 mg, 0.087 mmol), and glutaric anhydride (199 mg, 1.74 mmol) at 23 ºC. After 15 h, the reaction mixture was quenched with 1N $HCl_{(aq)}$ (3 mL), concentrated, and purified by column chromatography (2–4% MeOH/$CH_2Cl_2$ (0.1% AcOH)) to provide **5** (292 mg, 82% over 2 steps). $^1$H NMR (400 MHz, DMSO-$d_6$) δ 12.08 (bs, 1H), 10.08 (s, 1H), 7.69 (s, 4H), 7.15 (s, 1H), 6.01 (m, 1H), 5.40 (dq, $J$ = 17.1, 1.6 Hz, 1H), 5.28 (dq, $J$ = 10.6, 1.3 Hz, 1H), 4.77 (s, 2H), 2.39 (t, $J$ = 7.5 Hz, 2H), 2.30 (t, $J$ = 7.5 Hz, 2H), 1.84 (q, $J$ = 7.0 Hz, 2H); $^{13}$C NMR (101 MHz, DMSO-$d_6$) δ 174.68, 171.40, 162.36, 153.94, 153.78, 140.11, 132.73, 124.94, 123.93, 119.64, 118.59, 118.06, 105.03, 65.38, 55.30, 45.62, 33.44, 20.83.

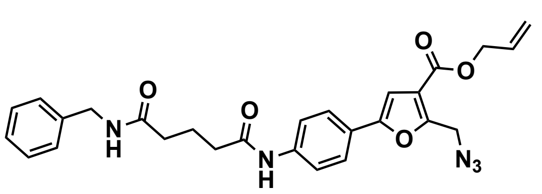

**6a**

*Allyl 2-(azidomethyl)-5-(4-(5-(benzylamino)-5-oxo pentanamido)phenyl)furan-3-carboxylate* (**6a**). **5** (50 mg, 0.12 mmol) was diluted in DMF (0.03 M, 4 mL) and treated with DIPEA (105 µL, 0.60 mmol) and HATU (57 mg, 0.15 mmol) at 23 °C. After 15 min, the reaction mixture was treated with benzylamine (16 μL, 0.15 mmol) and stirred for an additional 18 h. The reaction mixture was concentrated, diluted in $CH_2Cl_2$ (100 mL), and washed with $NaHCO_{3(sat)}$ (100 mL), $H_2O$ (100 mL), and brine (100 mL). The isolated organic layer was dried over $Na_2SO_4$ and concentrated to dryness to provide the crude product that was used without further purification. $^1$H NMR (400 MHz, $CDCl_3$) δ 8.89 (bs, 1H), 7.63 (d, $J$ = 8.6 Hz, 2H), 7.57 (d, $J$ = 8.6 Hz, 2H), 7.27 (m, 5H), 6.87 (s, 1H), 6.55 (bs, 1H), 6.03 (m, 1H), 5.41 (dq, $J$ = 17.3, 1.5 Hz, 1H), 5.33 (dq, $J$ = 10.3, 1.4 Hz, 1H), 4.80 (dt, $J$ = 5.8 Hz, 2H), 4.70 (s, 2H), 4.41 (d, $J$ = 5.6 Hz, 2H), 2.47 (t, $J$ = 6.9 Hz, 2H), 2.37 (t, $J$ = 7.0 Hz, 2H), 2.05 (q, $J$ = 7.0 Hz, 2H); $^{13}$C NMR (101 MHz, $CDCl_3$) δ 172.89, 171.43, 162.63, 153.98, 153.37, 138.67, 138.14, 131.84, 128.71, 127.62, 127.50, 124.79, 119.93, 118.84, 118.16, 104.70, 65.48, 45.70, 43.57, 36.37, 35.12, 21.93.

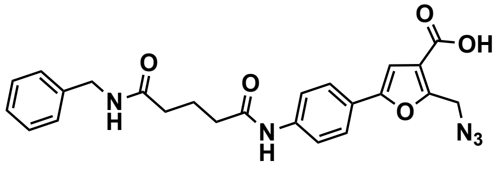


**7a**

*2-(Azidomethyl)-5-(4-(5-(benzylamino)-5-oxopentanamido)phenyl)furan-3-carboxylic acid* (**7a**). Crude **6a** was diluted in DMF (0.06 M, 2 mL) and treated with pyrrolidine (22 μL, 0.26 mmol) and $Pd(PPh_3)_4$ (7 mg, 0.006 mmol) at 23 °C. After 1.5 h, the reaction mixture was concentrated to dryness abd purified by column chromatography (2–6% MeOH/$CH_2Cl_2$ (0.1% AcOH)) to provide **7a** (24 mg, 43% over 2 steps). $^1$H NMR (400 MHz, DMSO-$d_6$) δ 13.17 (bs, 1H), 10.08 (s, 1H), 8.35 (t, $J$ = 6.0 Hz), 7.70 (s, 4H), 7.32 (t, $J$ = 7.5 Hz, 2H), 7.25 (t, $J$ = 6.3 Hz, 3H), 7.11 (s, 1H), 4.79 (s, 2H), 4.29 (d, $J$ = 5.9 Hz, 2H), 2.37 (t, $J$ = 7.4 Hz, 2H), 2.23 (t, $J$ = 7.7 Hz, 2H), 1.87 (q, $J$ = 7.5 Hz, 2H); $^{13}$C NMR (101 MHz, DMSO-$d_6$) δ 172.15, 171.53, 164.41, 153.58, 153.18, 140.13, 128.73, 127.65, 127.16, 124.90, 124.20, 119.71, 105.70, 50.72, 45.57, 42.49, 36.23, 35.03, 21.64.

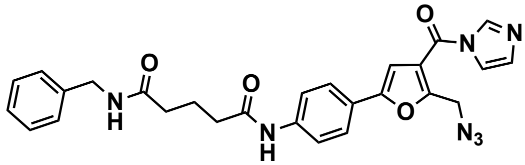


**8a**

*N1-(4-(5-(Azidomethyl)-4-(1H-imidazole-1-carbonyl)furan-2-yl)phenyl)-N5-benzylglutaramide* (**8a**). **7a** (20 mg, 0.043 mmol) was diluted in DMSO (0.02 M, 2 mL) and treated with 1,1′-carbonyldiimidazole (CDI) (14 mg, 0.086 mmol) at 23 °C. After 3.5 h, the reaction was quenched with $H_2O$ (3 mL) and concentrated via lyophilization. The crude reside was purified by reverse-phase semi-preparative

HPLC method B to provide **8a** (16 mg, 76%). $^1$H NMR (400 MHz, DMSO-$d_6$) δ 10.10 (s, 1H), 8.43 (t, *J* = 1.0 Hz, 1H), 8.34 (t, *J* = 6.0 Hz, 1H), 7.81 (t, *J* = 1.5 Hz, 1H), 7.76 (d, *J* = 8.8 Hz, 2H), 7.73 (d, *J* = 8.8 Hz, 2H), 7.40 (s, 1H), 7.32 (m, 2H), 7.25 (m, 3H), 7.19 (dd, *J* = 1.7, 0.8 Hz, 1H), 4.76 (s, 2H), 4.28 (d, *J* = 5.9 Hz, 2H), 2.38 (t, *J* = 7.0 Hz, 2H), 2.23 (t, *J* = 7.4 Hz, 2H), 1.87 (q, *J* = 7.6 Hz, 2H); LRMS (ESI$^+$) 512.19 $[M+H]^+$.

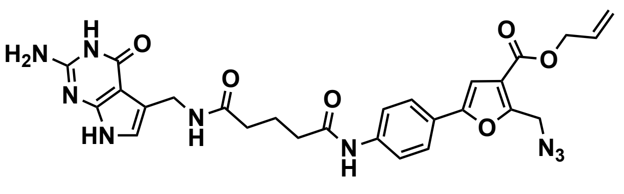

6b

*Allyl 5-(4-(5-(((2-amino-4-oxo-4,7-dihydro-3H-pyrrolo[2,3-d]pyrimidin-5-yl)methyl) amino)-5-oxopentanamido)phenyl)-2-(azido methyl)furan-3-carboxylate* (**6b**). **5** (24 mg, 0.058 mmol) was diluted in DMF (0.03 M, 2 mL) and treated with DIPEA (34 µL, 0.19 mmol) and HATU (22 mg, 0.058 mmol) at 23 °C. After 15 min, preQ1 hydrochloride (12 mg, 0.048 mmol) was added and the reaction was stirred for an additional 24 h. The reaction mixture was concentrated to dryness and purified by column chromatography (6–8% MeOH/$CH_2Cl_2$) to provide **6b** (9 mg, 33%).$^1$H NMR (400 MHz, DMSO-$d_6$) δ 10.82 (s, 1H), 10.37 (s, 1H), 10.06 (s, 1H), 8.21 (t, *J* = 5.5 Hz, 1H), 7.71 (d, *J* = 1.7 Hz, 4H), 7.20 (s, 1H), 6.50 (t, *J* = 1.1 Hz, 1H), 6.10 (s, 2H), 6.04 (m, 1H), 5.42 (dq, *J* = 17.2, 1.7 Hz, 1H), 5.30 (dq, *J* = 10.5 Hz, 1H), 4.81 (s, 2H), 4.79 (dt, *J* = 5.4, 1.5 Hz, 1H), 4.30 (d, *J* = 4.9 Hz, 2H), 2.35 (t, *J* = 7.3 Hz, 2H), 2.17 (t, *J* = 7.3 Hz, 2H), 1.84 (q, *J* = 7.3 Hz, 2H).

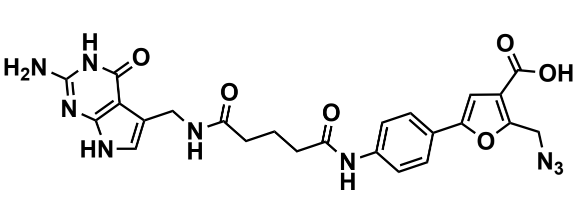

7b

*5-(4-(5-(((2-amino-4-oxo-4,7-dihydro-3H-pyrrolo [2,3-d]pyrimidin-5-yl)methyl)amino)-5-oxopentan amido)phenyl)-2-(azidomethyl)furan-3-carboxylic acid* (**7b**). **6b** (9 mg, 0.016 mmol) was diluted in DMF (0.02 M, 1 mL) and treated with pyrrolidine (3 µL, 0.033 mmol) and $Pd(PPh_3)_4$ (1 mg, $0.75\times10^{-3}$ mmol) at 23 °C. After 2 h, the reaction mixture was concentrated to dryness and purified by reverse-phase semi-preparative HPLC with method A to provide **7b** (5.5 mg, 68%).$^1$H NMR (400 MHz, DMSO-$d_6$) δ 10.82 (s, 1H), 10.44 (bs, 1H),

10.04 (s, 1H), 8.22 (t, $J$ = 5.5 Hz, 1H), 7.68 (d, $J$ = 2.0 Hz, 4H), 7.05 (s, 1H), 6.49 (t, $J$ = 1.1 Hz, 1H), 6.14 (s, 2H), 4.80 (s, 2H), 4.30 (d, $J$ = 5.4 Hz, 2H), 2.34 (t, $J$ = 7.4 Hz, 2H), 2.17 (t, $J$ = 7.4 Hz, 2H), 1.84 (q, $J$ = 7.2 Hz, 2H).

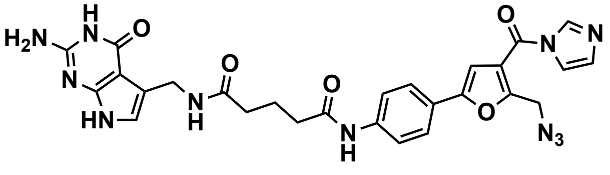

8b

*N1-((2-amino-4-oxo-4,7-dihydro-3H-pyrrolo [2,3-d]pyrimidin-5-yl)methyl)-N5-(4-(5-(azido methyl)-4-(1H-imidazole-1-carbonyl)furan-2-yl)phenyl)glutaramide* (**8b**). **7b** (5 mg, 0.0093 mmol) was diluted in DMSO (0.01 M, 1 mL) and treated with CDI (3 mg, 0.018 mmol) at 23 °C. After 24 h, the reaction mixture was quenched with $H_2O$ (1 mL), concentrated via lyophilization, and purified by reverse-phase semi-preparative HPLC with method B to provide probe **8b** (3.6 mg, 41%). $^1$H NMR (400 MHz, DMSO-$d_6$) δ 10.82 (s, 1H), 10.35 (s, 1H), 10.08 (s, 1H), 8.43 (t, $J$ = 1.0 Hz, 1H), 8.21 (t, $J$ = 5.3 Hz, 1H), 7.81 (t, $J$ = 1.4 Hz, 1H), 7.77 (d, $J$ = 8.9 Hz, 2H), 7.72 (d, $J$ = 9.0 Hz, 2H), 7.40 (s, 1H), 7.19 (dd, $J$ = 1.7, 0.8 Hz, 1H), 6.50 (t, $J$ = 1.1 Hz, 1H), 6.09 (s, 2H), 4.76 (s, 2H), 4.30 (d, $J$ = 5.4 Hz, 2H), 2.36 (t, $J$ = 7.8 Hz, 2H), 2.18 (t, $J$ = 7.2 H, 2H), 1.84 (q, $J$ = 7.2 Hz, 2H) ; LRMS (ESI+) 548.19[M+H] (observed mass of converted active probe into methyl ester after treatment with methanol).

### D. NMR Spectra

#### Compound 2

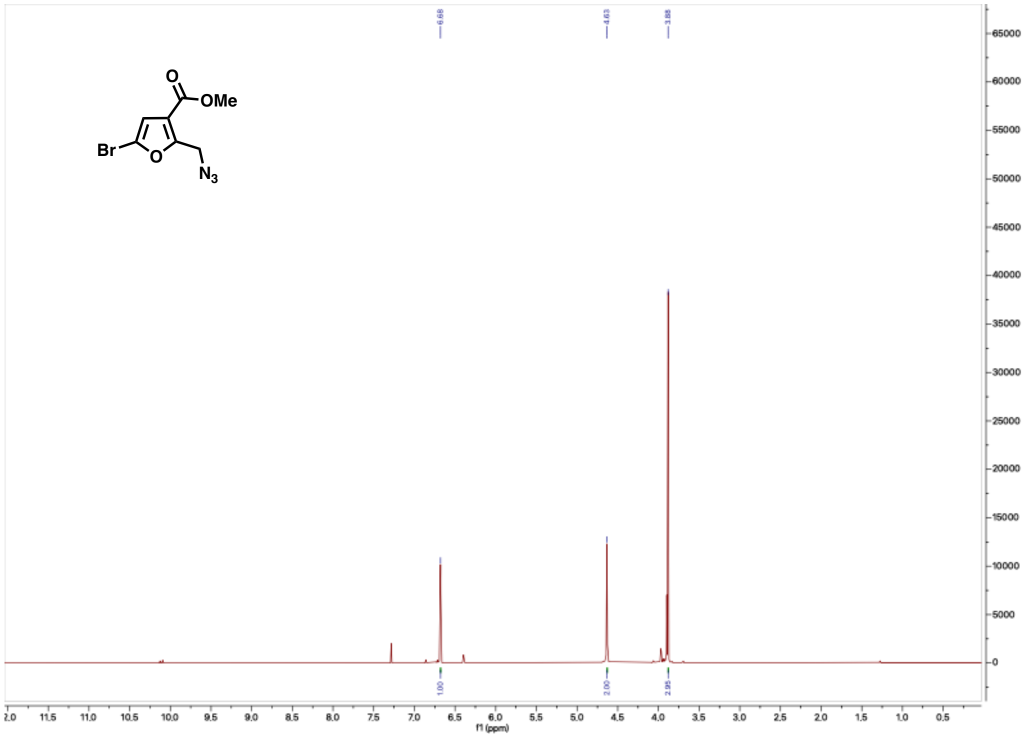


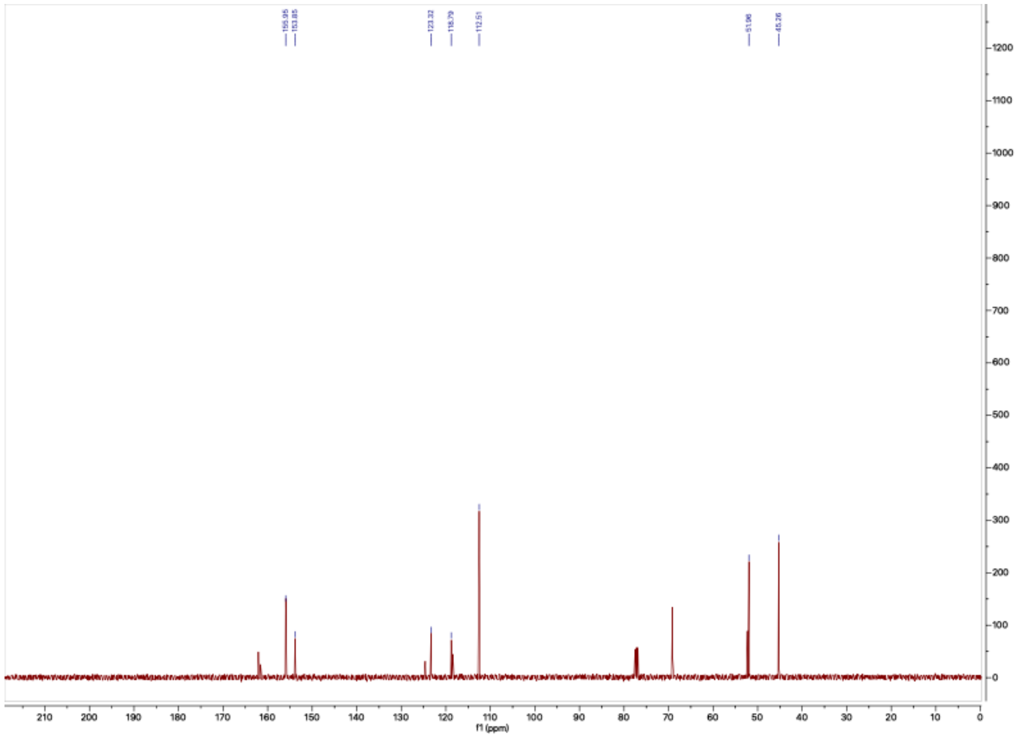

### Compound 3

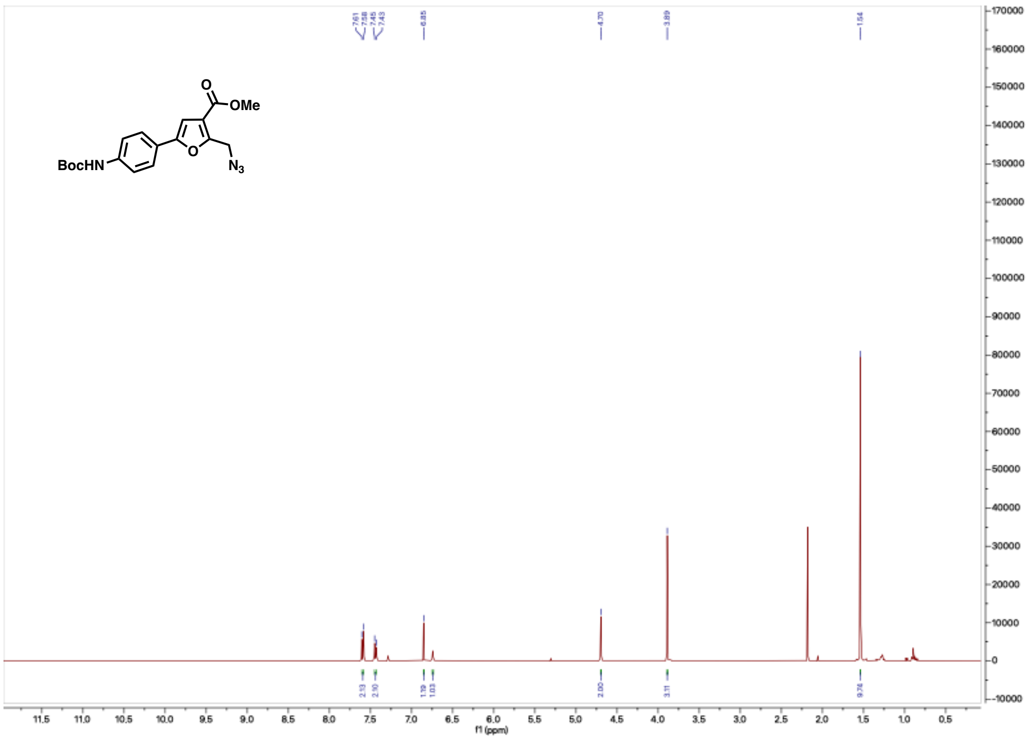


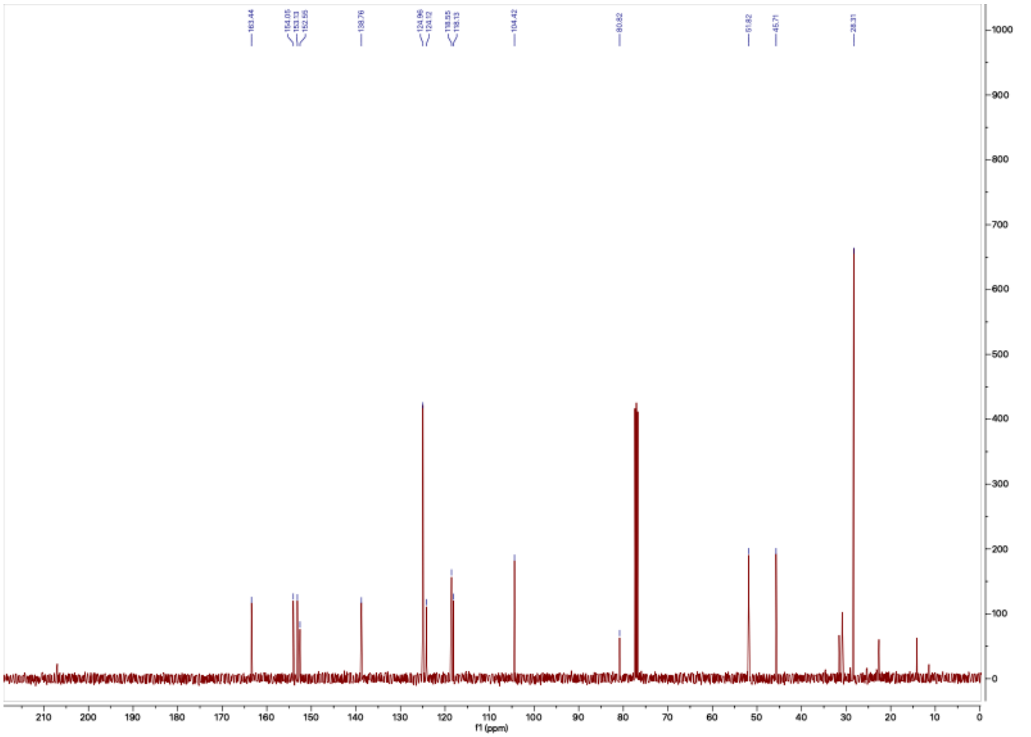

**Compound 4**

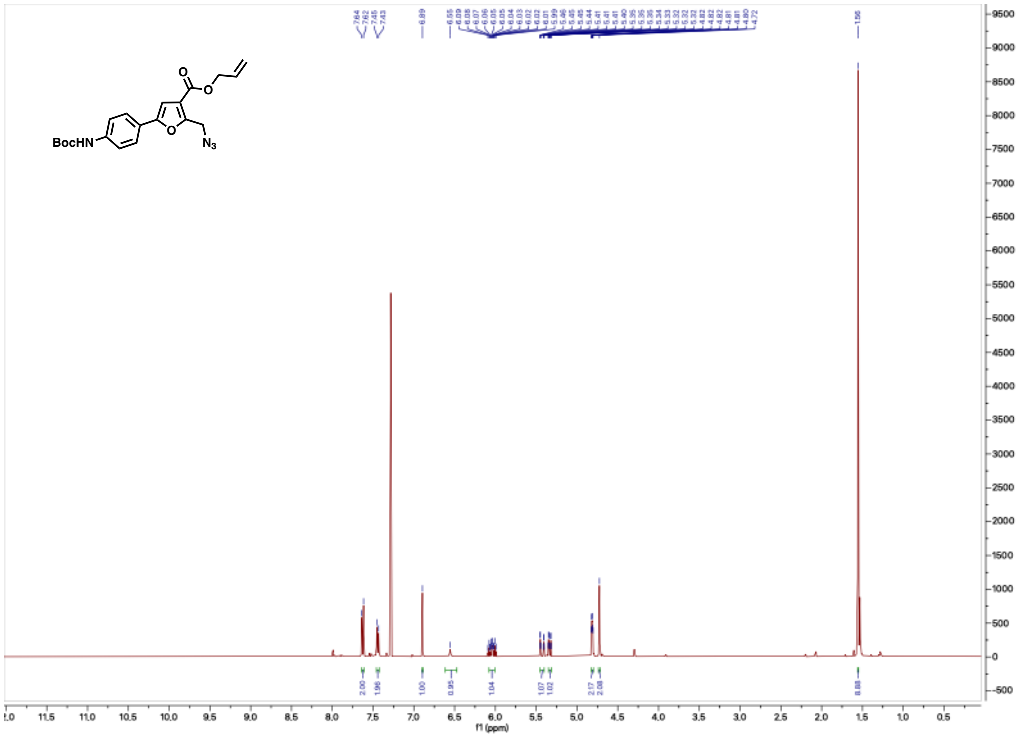


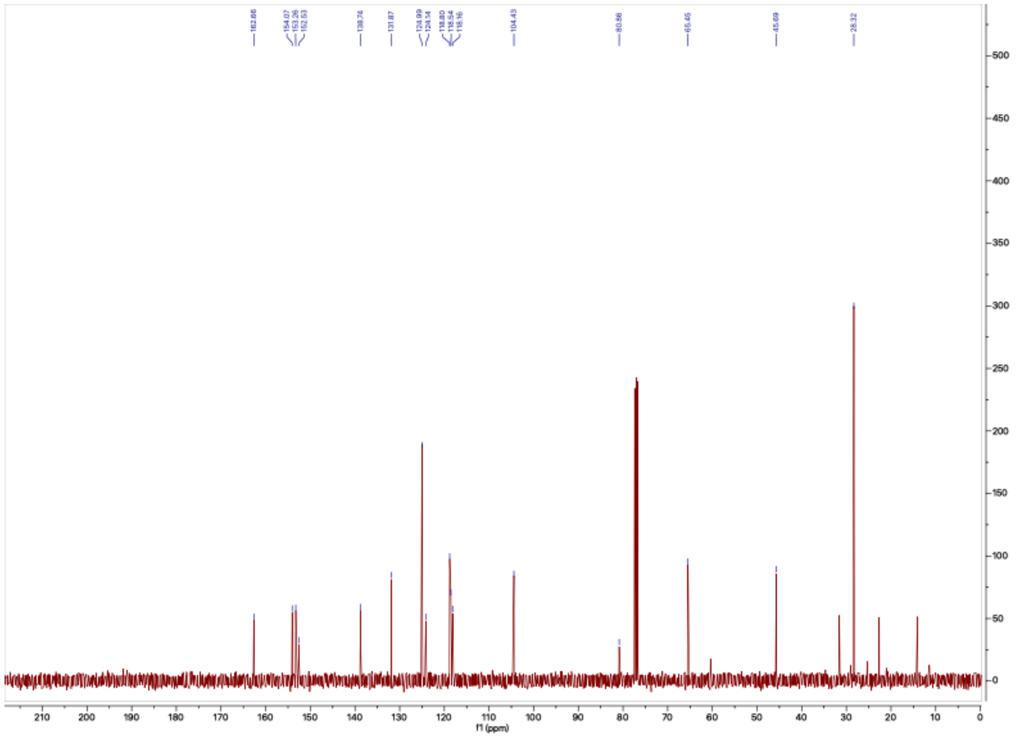

**Compound 5**

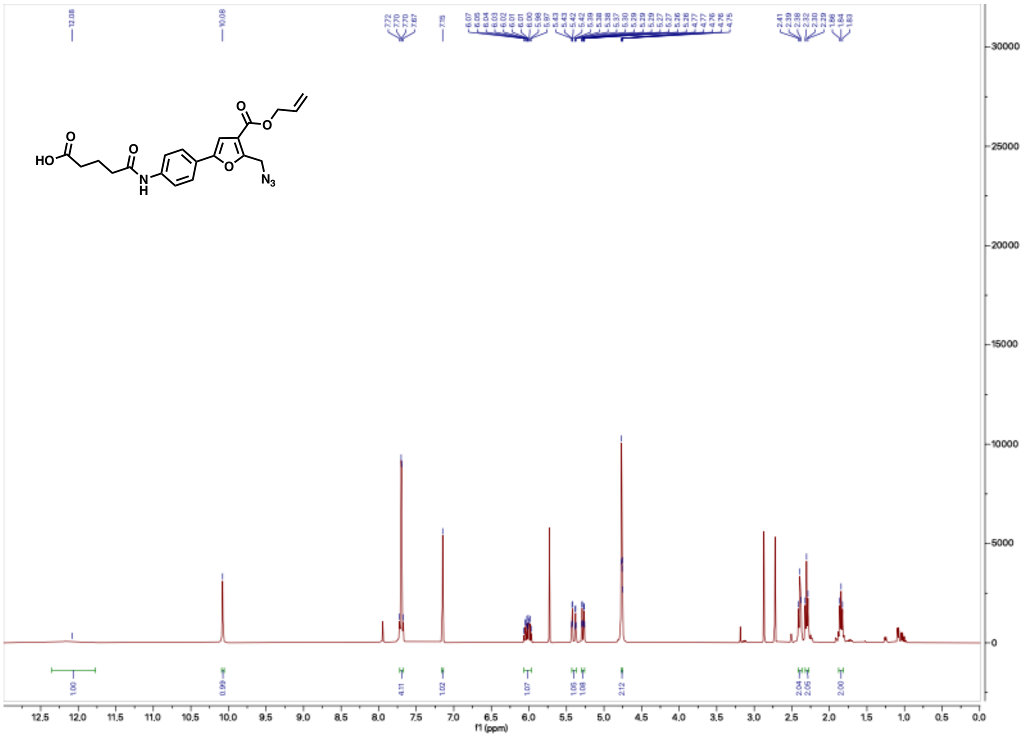

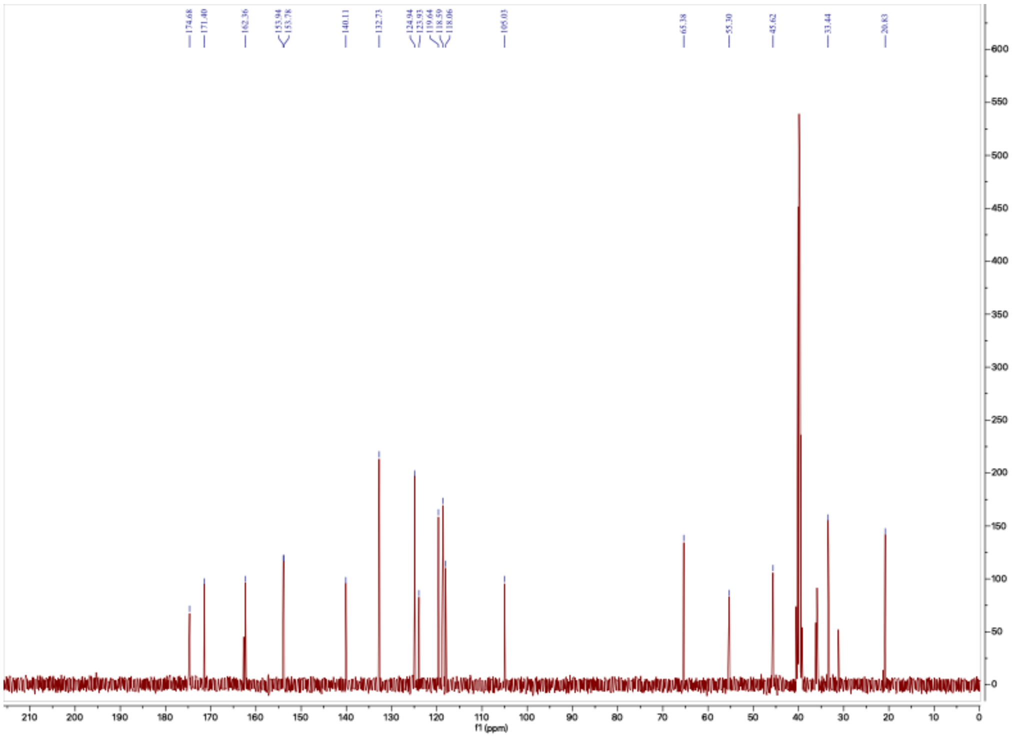

#### Compound 6a

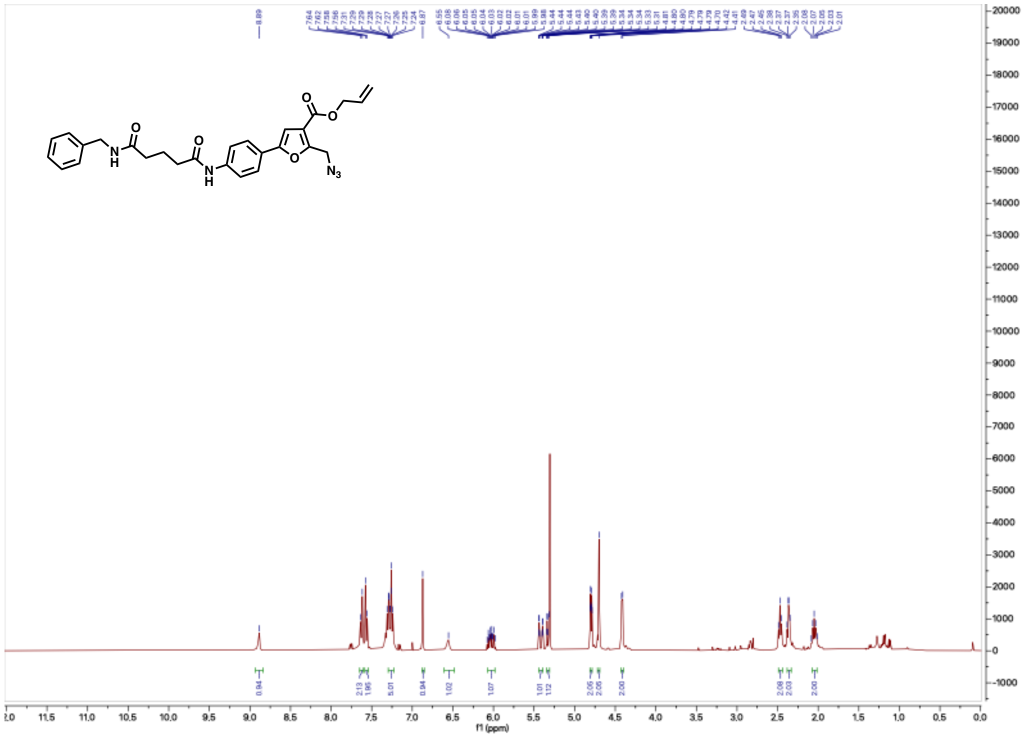

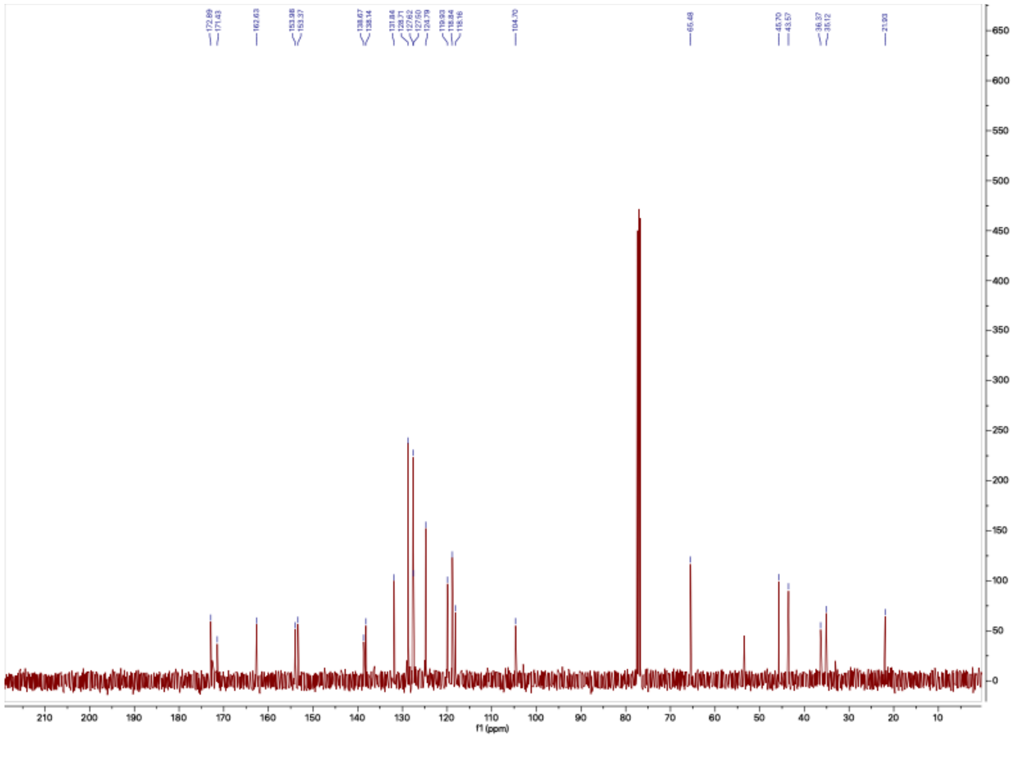

#### Compound 7a

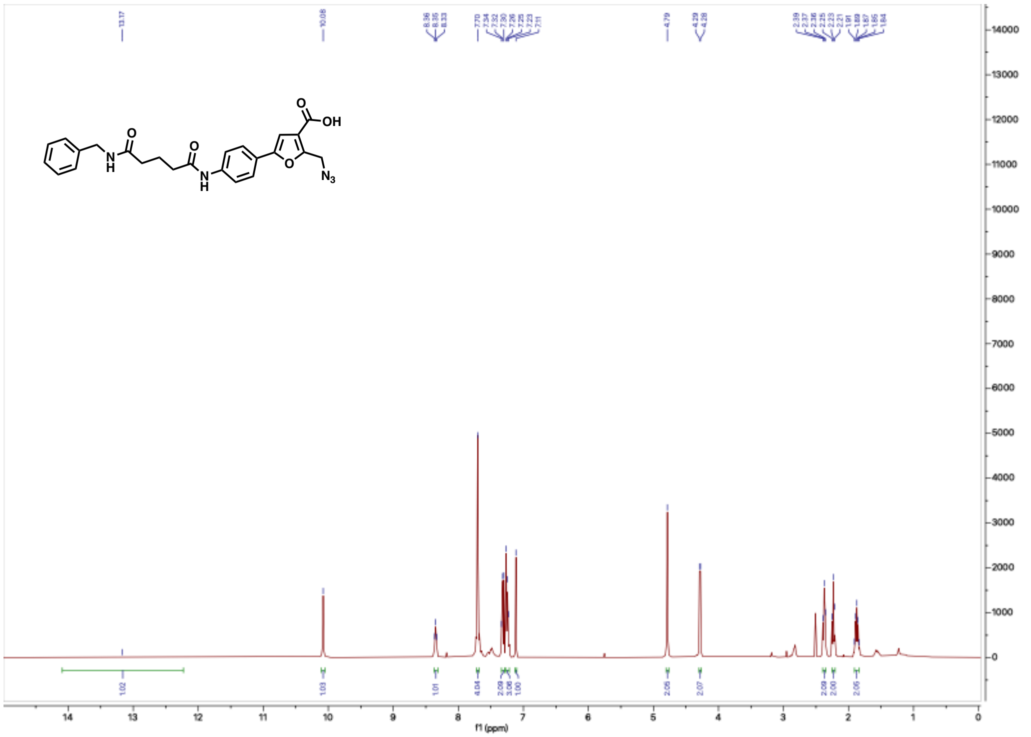

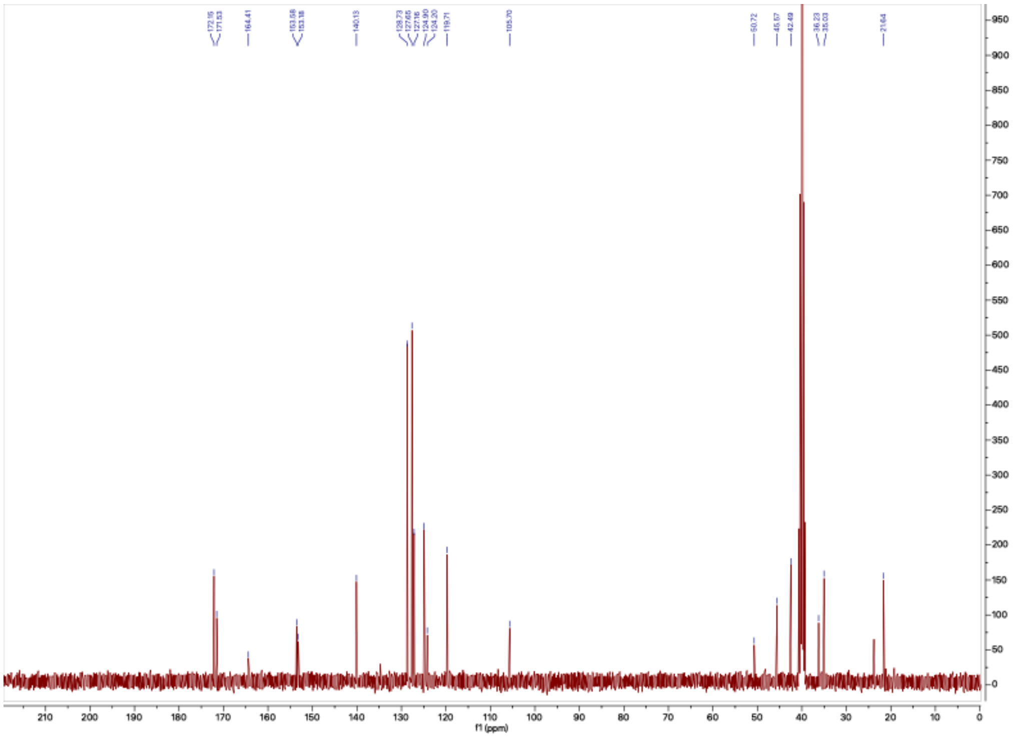

**Compound 8a**

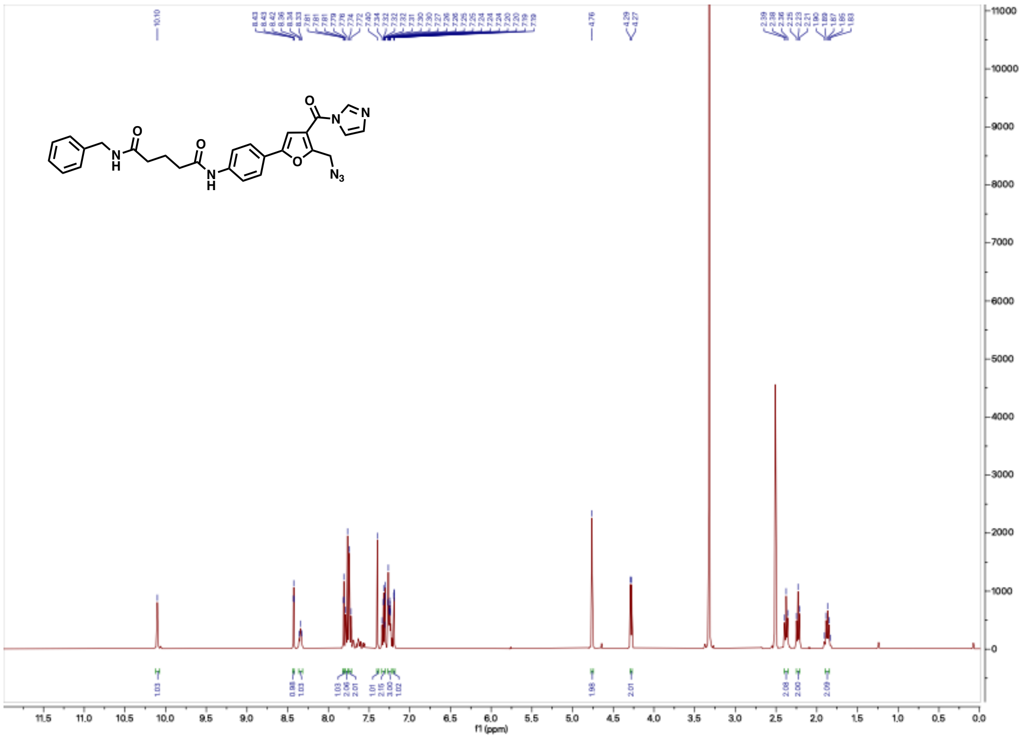

#### Compound 6b

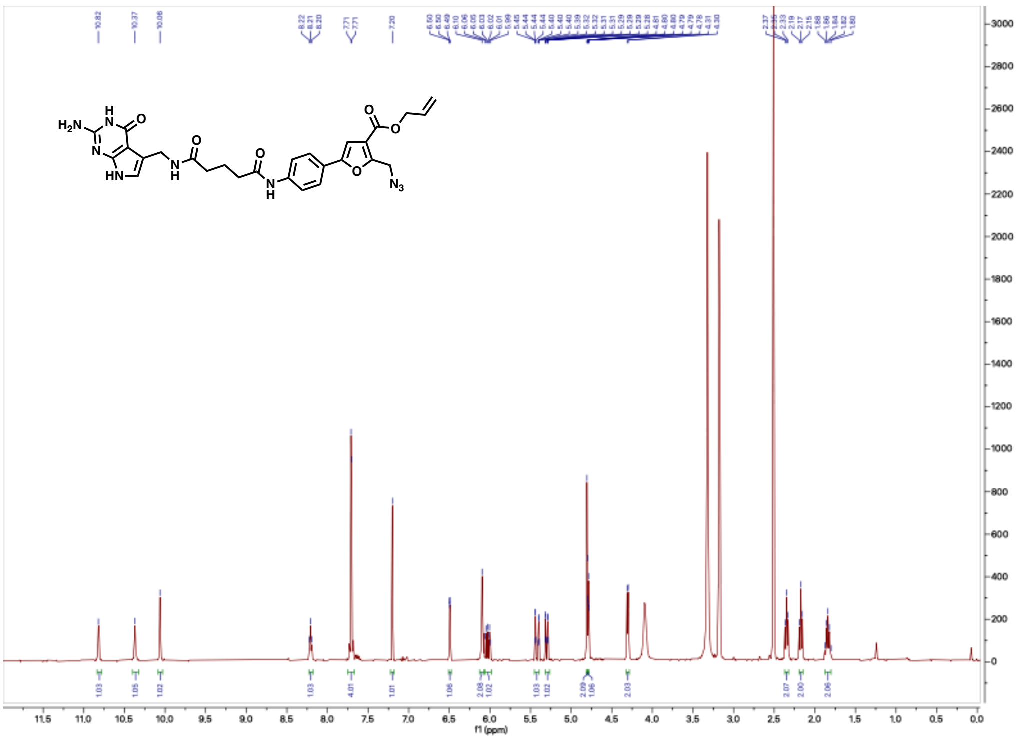

#### Compound 7b

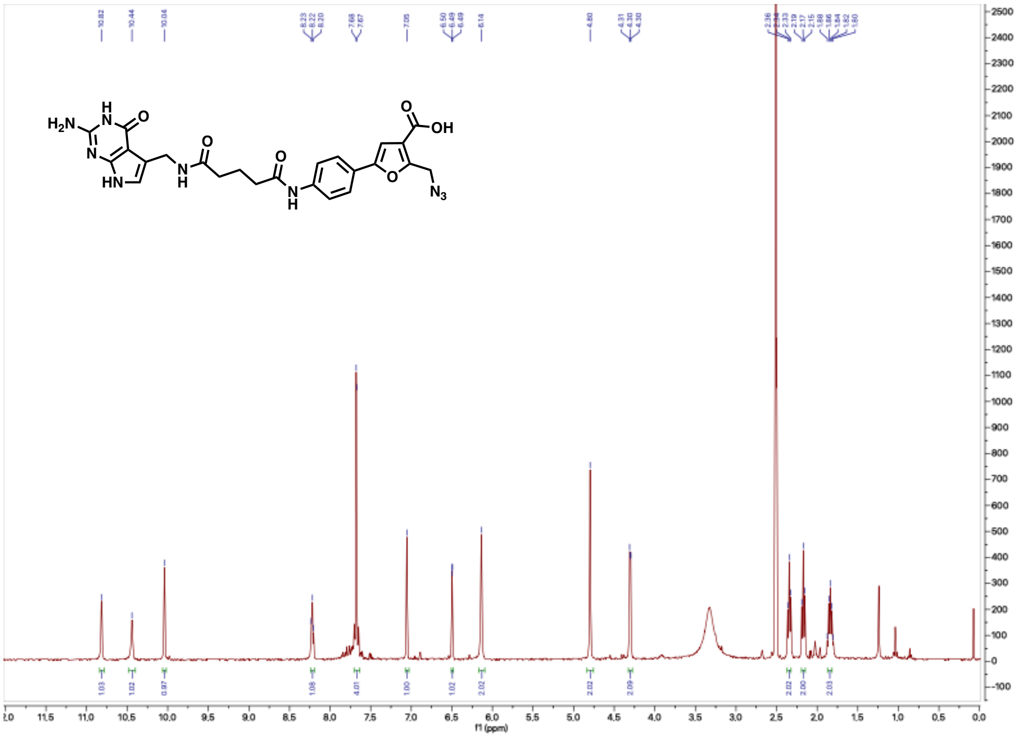

#### Compound 8b

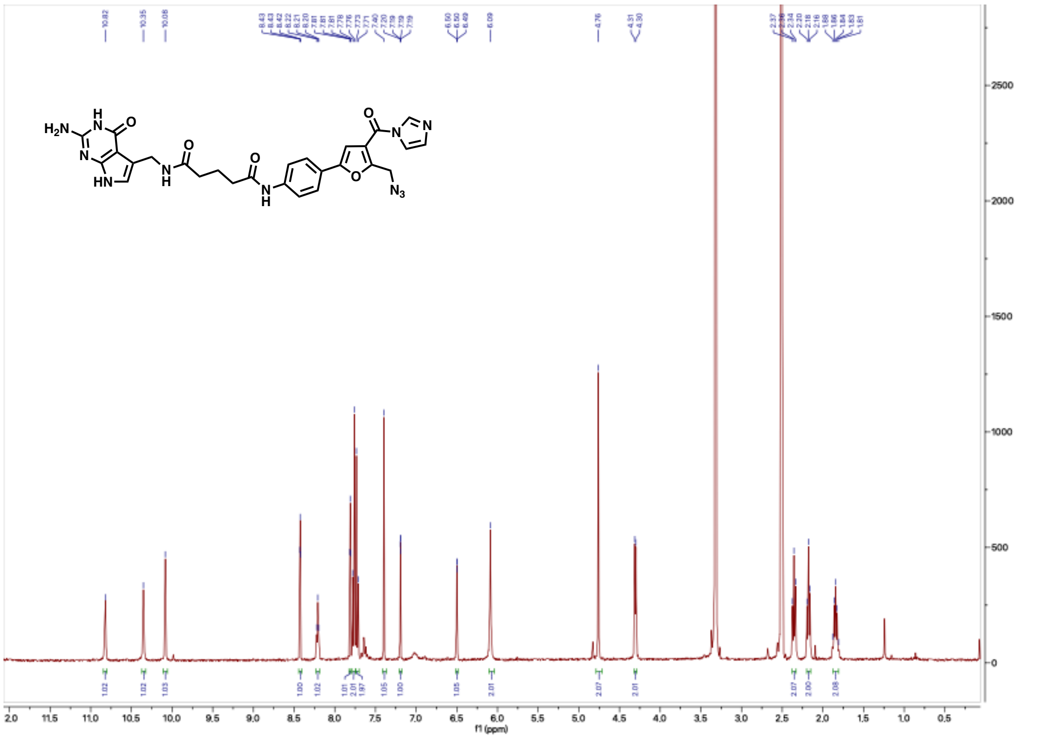
